# A forskolin-mediated increase in cAMP promotes T helper cell differentiation into the Th1 and Th2 subsets rather than into the Th17 subset

**DOI:** 10.1101/2022.06.26.497628

**Authors:** Petra Daďová, Antónia Mikulová, Radim Jaroušek, Stjepan Uldrijan, Lukáš Kubala

## Abstract

The cyclic adenosine monophosphate (cAMP) signaling pathway is involved in various physiological and pathophysiological processes. Forskolin (FSK), a labdane diterpene well known as an activator of cAMP production, is suggested to possess significant immunomodulatory potential. However, the specific effects of elevated cAMP levels caused by the FSK-mediated activation of adenylate cyclase (AC) on T helper (Th) cell differentiation and functions are still unclear. We speculated that the increased levels of cAMP in Th cells affect the differentiation program of distinct Th populations differently, in particular the development of Th1, Th2, and Th17 subsets. Only minor changes in the expressions of isoforms of ACs and phosphodiesterases (PDE), enzymes responsible for the degradation of cAMP, were observed in differentiating human Th cells with prevailing *ADCY1, ADCY3, ADCY7*, and *ADCY9* and *PDE3B*, *PDE4A/B/D*, *PDE7A/B*, and *PDE8A* isoforms. FSK mediated elevation in Th1-specific markers reinforcing the Th1 cell phenotype. The differentiation of Th2 was not altered by FSK, though cell metabolism was affected. In contrast, the Th17 immunophenotype was severely suppressed leading to the highly specific upregulation of the level of CXCL13. The causality between FSK-elicited cAMP production and the observed modulation of Th2 differentiation was proven by using cAMP inhibitor 2’,5’-dideoxyadenosine that reverted the FSK effects. Overall, an FSK-mediated cAMP increase has an effect on Th1, Th2 and Th17 differentiation and can contribute to the identification of novel therapeutic targets for the treatment of Th cell-related pathological processes.

## 1. Introduction

Cyclic adenosine monophosphate (cAMP), as an important secondary messenger of cell signaling, is involved in various physiological and pathophysiological processes, including embryogenesis, cardiac function, pain reception, learning, and aging. The synthesis of cAMP in the cell is catalyzed by adenylate cyclases (AC). Many clinically used drugs act as ligands of G-protein coupled receptors (GPCRs) that directly modulate ACs or target the cAMP-degrading enzymes phosphodiesterases (PDEs). To date, 10 mammalian AC isoforms (AC1-10) that differ in the ways their enzymatic activity is regulated as well as in their tissue distributions have been identified. In immune cells, three dominant AC isoforms were detected (AC3, AC7 and AC9) (***Dessauer et al., 2017***). Forskolin (FSK), the labdane diterpene isolated from the root of the herb *Coleus Forskohlii (**Mitra et al., 2020**)*, is well known as a non-specific activator of all membrane-bound ACs. In contrast to the function of ACs, the intracellular concentration of cAMP is regulated by PDE, which belongs to the large family of enzymes with the main function of degrading cAMP and converting it into inactive 5’-AMP. cAMP in cells regulates effector molecules, including protein kinase A (PKA), Epac, and cyclic-nucleotide gated channels.

It is well known that the cAMP pathway plays a crucial role in the management of immune cell signaling, including that of T helper (Th) cells. Th cells, after being activated and differentiated into distinct effector subtypes, mediate the immune response and communicate with the target cells (e.g. T cytotoxic cells, B cells, or macrophages) by secreting specific cytokines. Upon activation, Th-naïve cells are polarized by cytokines into specific Th subsets, including Th1, Th2, and Th17, that carry out specific effector functions. Differentiation into the Th1 subset is driven by interleukin (IL)-12 in response to an intracellular pathogen. The T-box transcription factor (T-bet), encoded by the *TBX21* gene, is the master regulator of Th1 differentiation and induces the production of the Th1 effector cytokine, interferon (IFN)-γ. IL-4 is both a polarizing and an effector cytokine of the Th2 subset, which drives the positive feedback machinery of Th2 lineage commitment (***Zhu, 2010***). Th2-generated cytokines, including IL-4, IL-5, and IL-13, promote B cells to produce immunoglobulin (Ig) E and IgA. The main Th2-specific transcription factor Gata-binding protein (GATA3) is involved in enhancing Th2 cytokine production, selective proliferation, as well as the inhibition of Th1 differentiation. Innate immune cells produce large amounts of both IL-6 and tumor necrosis factor (TGF) β in response to extracellular bacteria and fungi. These cytokines together with IL-23 and IL-1β create the environment for successful Th17 differentiation. Retinoic acid receptor-related orphan receptor gamma-T (RORγt), encoded by the *RORC2* gene, is the main transcription factor of Th17 cells, also characterized by its ability to produce effector cytokines such as IL-17A and IL-17F (***Korn et al., 2009***). TGF-β is also essential for the induction of Th-naïve cell differentiation into FoxP3+ regulatory T cells (Treg). This results in a close relationship with the Th17 subset, and suggests the possibility of Th17/Treg plasticity, even though both Th subsets play opposite roles in inflammatory processes (***Sehrawat and Rouse, 2017***). In general, a disbalance of Th-subsets in the body is connected with the altered regulation of both specific and innate immune system mechanisms and consequently to the development of a pathological status. As an example, pathogenically dominant Th1 was associated with autoimmune disease, like type I diabetes, Th2 with allergy and asthma, and Th17 with Crohn’s disease and lupus ***(Korn et al., 2009, Pennock et al., 2013)***.

The importance of AC in the proper function of the immune system can be suggested by the consequences of mutations in genes coding some ACs (*ADCY* genes) in individuals suffering from autoimmune disorders, including Crohn’s disease and ulcerative colitis. Even though these pathologies are considered multifactorial, this observation supports evidence of a connection between AC activity and clinical significance. For example, a mutation in *ADCY3* (rs6545800) was identified in patients with inflammatory bowel diseases ***(Hulur et al., 2015, Jostins et al., 2012)***. A mutation in *ADCY7* (rs77150043) was suggested to play a role in the early onset of psoriasis and Crohn’s disease (***Li et al., 2015***), while another mutation in *ADCY7* (rs78534766) doubles the risk of the development of ulcerative colitis (***Luo et al., 2017***), all such mutations related to the deregulation of proper Th cell subpopulation balance.

For a long time, cAMP was understood to act as a potent suppressor of all immune cells, including Th cells. It was assumed that cAMP inhibits almost all downstream pathways of TCR stimulation and suppresses Th cell activation and proliferation, and cytokine production (***Arumugham and Baldari, 2017***). However, previous studies have revealed cAMP to play a more complex and diverse role in the activation and differentiation of diverse subpopulations of Th cells, particularly in the context of cAMP signaling pathway activation through the activation of GPCRs by their ligands such as prostaglandins (PG) or histamine. The potentiation of Th1 differentiation was reported after stimulation with PGE2 and PGI2 ***(Nakajima et al., 2010, Sakata et al., 2010, Yao et al., 2013, Yao et al., 2009)***. On the other hand, the elevated secretion of Th2 cytokines and the inhibited secretion of Th1 cytokines was induced by histamine ***(Elliott et al., 2001, Okamoto et al., 2009***). The promotion of Th2 differentiation and the inhibition of IL-2 and IFN-γ production was mediated (***Tokoyoda et al., 2004***). Th17 cell expansion was potentiated by PGE2-EP4 ***(Boniface et al., 2009, Sakata et al., 2010, Yao et al., 2009)***. In contrast, the inhibition of PDE activity disrupting cAMP degradation resulted in the suppression of the production of the Th17-specific effector cytokine IL-17A (***Ma et al., 2008, Schafer et al., 2014)***. Interestingly, the inhibition of PDE enhanced the development of Treg cells, shifting the Th17/Treg transdifferentiation program in the Treg direction (***Schafer et al., 2014***). Overall, current studies point to the importance of cAMP in the process of Th cell differentiation into particular subsets, suggesting the possible promotion of Th2 over Th1 differentiation and Treg over Th17. However, there is heterogeneity in the reported data that can be related to the diversity of complex intracellular signaling nets induced by different GPCR ligands. The importance of cAMP for Th cell differentiation in response to the direct and highly specific activation of AC by FSK, the most direct pathway to elevating the intracellular cAMP concentration, is yet to be established. The modulation of Th cell differentiation by FSK-mediated cAMP elevation could contribute to the identification of novel therapeutic targets for the treatment of Th cell-related pathological processes.

With the aim of describing the role of cAMP in the differentiation of mature human Th cells, a complex study of the FSK-mediated modulation of the differentiation of distinct Th subsets – Th1, Th2, and Th17 – was performed. In addition to the determination of typical transcription factors and effector cytokines, the differentiating Th subsets were also characterized on the basis of the expression of specific gene signatures using single-cell RNA sequencing (scRNAseq), specifying the cell immunophenotype in detail. A comprehensive analysis of the expressions of different isoforms of two key enzymes in the cAMP signaling pathway, ACs and PDEs, was also performed in differentiating Th cells.

## 2. Results

### Differentiation into Th1, Th2 and Th17 subsets

In our study, we worked with the three most abundant Th cell subsets in the human immune system – Th1, Th2 and Th17. First, Th naïve cells were activated with a combination of plate-coated anti-CD3 and soluble anti-CD28, followed by cell differentiation for 5 days in the presence of cocktails of specific cytokines and inhibitory antibodies for each subset (see Methods). Th cells stimulated with anti-CD3/anti-CD28 in the absence of cytokines and inhibitory antibodies served as an activated spontaneously differentiated control (ACT), whereas Th cells cultivated in the absence of both the activation stimulus and cytokines served as a resting cell control (NT ctrl). Apparently, not only activation but also a differentiation stimulus is essential for Th cell proliferation. The proliferation assay showed significant elevation of the number of activated and differentiated cells compared to NT ctrl. Moreover, the Th2 subset revealed a further significant increase compared to ACT, whereas Th1 and Th17 showed only mild elevation of cell numbers compared to ACT (**Supplementary data 1A**).

The targeted differentiation of activated Th cells into the Th1, Th2, and Th17 subsets was confirmed on the basis of the expressions of transcription factors (*TBX21*, *GATA3*, *RORC2*) and the production of effector cytokines (IFN-γ, IL-13, IL-17A) typical for particular Th subsets (**Supplementary data 2A-C, G-I**). As expected, in comparison to NT ctrl and ACT, Th1 revealed a significantly higher expression of *TBX21* and a significantly higher production of IFN-γ, Th2 revealed a significantly higher level of *GATA3* and a significantly higher production of IL-13, and Th17 revealed a significantly higher expression of *RORC2* and a significantly higher production of IL-17A.

Further, the selected samples underwent scRNAseq analysis, which provided detailed transcriptomic data for each subset (comprising approximately 700 single cells per subset; see Methods **Tab. 1**). scRNAseq allowed the identification of distinct Th subsets based on signature genes that are strongly connected to particular Th subsets, in addition to classic characterization based on transcription factors involved in the regulation of effector cytokine production. To combine the classic and the novel approaches, 1 transcription factor and 2 signature genes were chosen to characterize each Th cell subset. *TBX21*, *ANXA3* and *RUNX3* were found to be specific for Th1, *GATA3*, *LIMA1* and *MAOA* for Th2, and *RORC2*, *PALLD* and *IL23R* for Th17 (**Figure 1, Supplementary data 2D-F**).

**Figure 1.**
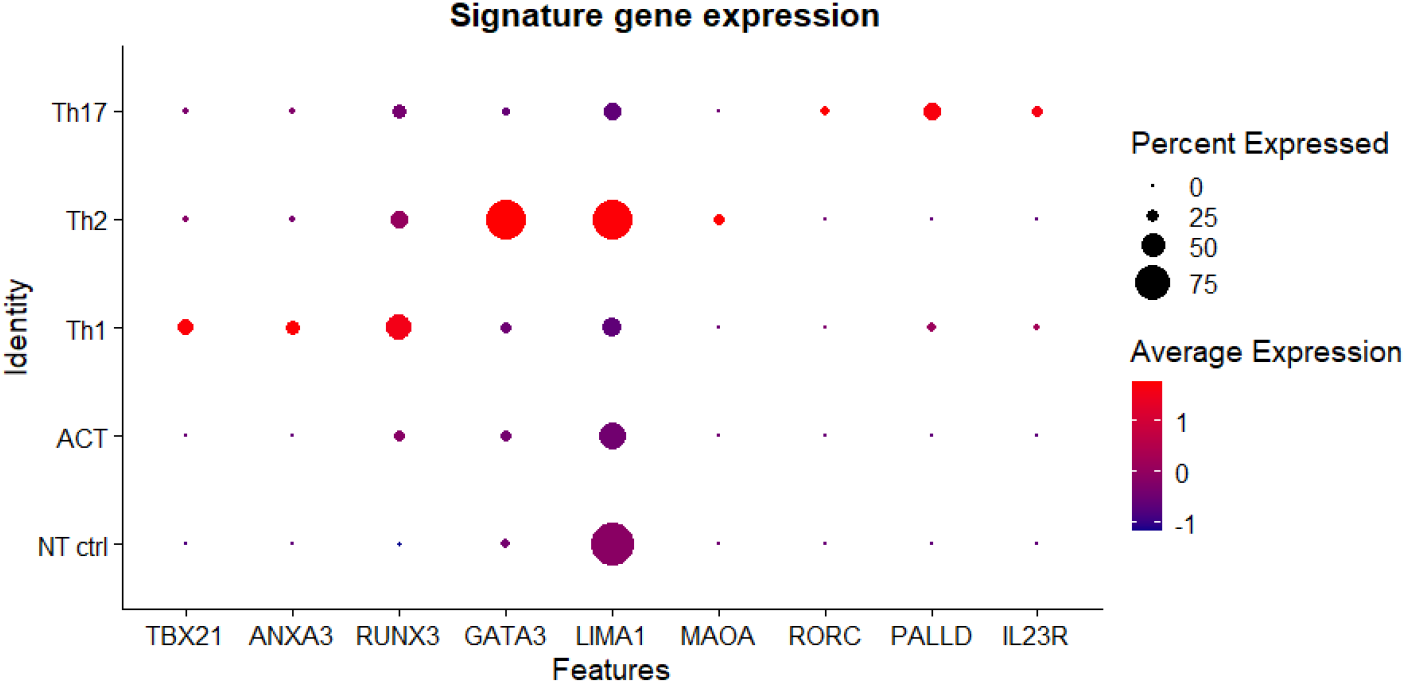
Gene expression of selected signature genes specific for Th1, Th2 or Th17 subset. DotPlot visualization of subset-specific genes selected for further analysis according to scRNAseq. Gene expressions of TBX21, ANXA3 and RUNX3 are specific to the Th1 subset, GATA3, LIMA1 and MAOA to the Th2 subset, and RORC, PALLD and IL23R to the Th17 subset. Different colors correspond to the scaled expression (Z-score); the dot size represents the percentage of positive cells.

### Analysis of the expressions of particular ACs and PDEs in Th cells

To understand the importance of the cAMP signalling pathway in Th cell differentiation, we used FSK as the main non-specific activator of all membrane-bound ACs. FSK (10 μM) was added to Th cell cultures during the differentiation process 20 h after activation, and, as expected, induced a significant increase in intracellular cAMP levels (**Supplementary data 3**). Interestingly, FSK did not significantly affect the potential of Th1 cells to proliferate, though proliferation upon FSK treatment was partially impaired in the case of Th2 and Th17 cells (**Supplementary data 1B**).

To clarify which ACs are responsible for cAMP production in particular Th differentiation lineages, the gene expressions of all membrane-bound AC isoforms (*ADCY1-9*), as well as the soluble one (ADCY10), were determined by RT-qPCR. Generally, four different AC isoforms, *ADCY1, ADCY3, ADCY7*, and *ADCY9*, were detectable in Th cells (**Figure 2**). The resting cells (NT ctrl) exhibited different AC isoform gene expression profiles compared to the activated and differentiated cells. *ADCY7* is the dominant isoform in resting cells, whereas *ADCY3* becomes dominant after cell activation. *ADCY1* expression was detected only in activated and differentiated cells, which was statistically significant in the case of ACT and the Th1, and Th2 subsets. *ADCY9* expression was revealed to be significantly higher in ACT and Th1 compared to NT ctrl. Surprisingly, there was no significant difference between AC expression profiles in the ACT, Th1, Th2, or Th17 subsets (**Figure 2**). The activation of ACs by FSK did not promote changes in the gene expression of AC isoforms (**Supplementary data 4**).

**Figure 2.**
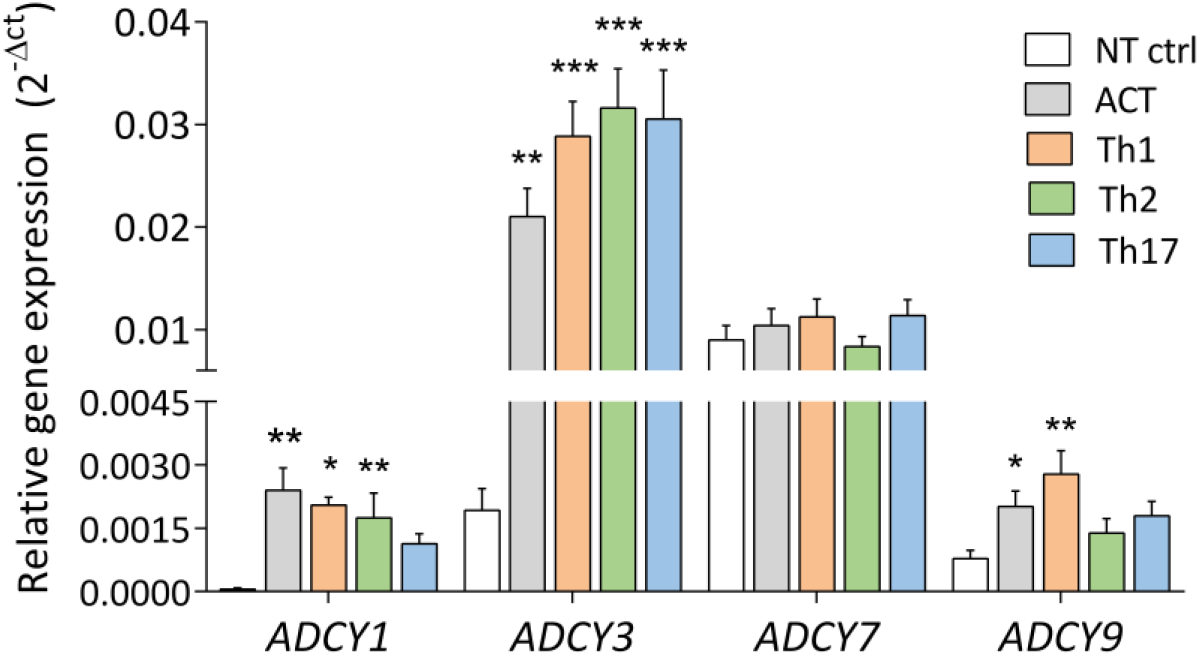
Analysis of adenylate cyclase (ADCY) isoform expression in Th1, Th2 and Th17 cells. Expression of detectable AC isoforms (ACDY1, ADCY3, ADCY7 and ADCY9) in the Th subsets (Th1, Th2 and Th17) and both non-treated (NT) and activated (ACT) controls. Statistical significance was examined by one-way ANOVA. Mean ± SEM, n=5-10, * p < 0.05, ** p < 0.01, *** p < 0.001 (vs NT ctrl)

The second kind of key enzymes regulating cAMP intracellular signaling are PDEs, which degrade cAMP to maintain homeostasis in the cell. The PDE family comprises a large number of genes; thus, the gene expression analysis was focused on the seven most abundant PDEs – namely, *PDE3B, PDE4A/B/D*, *PDE7A/B*, and *PDE8A* – that were revealed to be expressed in activated and differentiated Th cells by scRNAseq analysis (**Figure 3A, Supplementary data 5A**.), suggesting high variability among different Th cell subsets. RT-qPCR analysis identified *PDE3B* and *PDE7A* to have a higher expression in the NT ctrl compared to differentiated cells, except for *PDE3B* in ACT and Th1. In contrast, the isoforms *PDE4A* and *PDE7B* showed very low expression in NT ctrl (**Figure 3A**). Interestingly, *PDE4A* was absent in the Th17 subset and *PDE4B* was low in the Th2 subset. *PDE4B* and *PDE4D* were found to be specific to Th1. In addition, we found *PDE7B* to be specific to the Th2 subset. PDE7A and PDE8A did not differ among different Th subsets.

**Figure 3.**
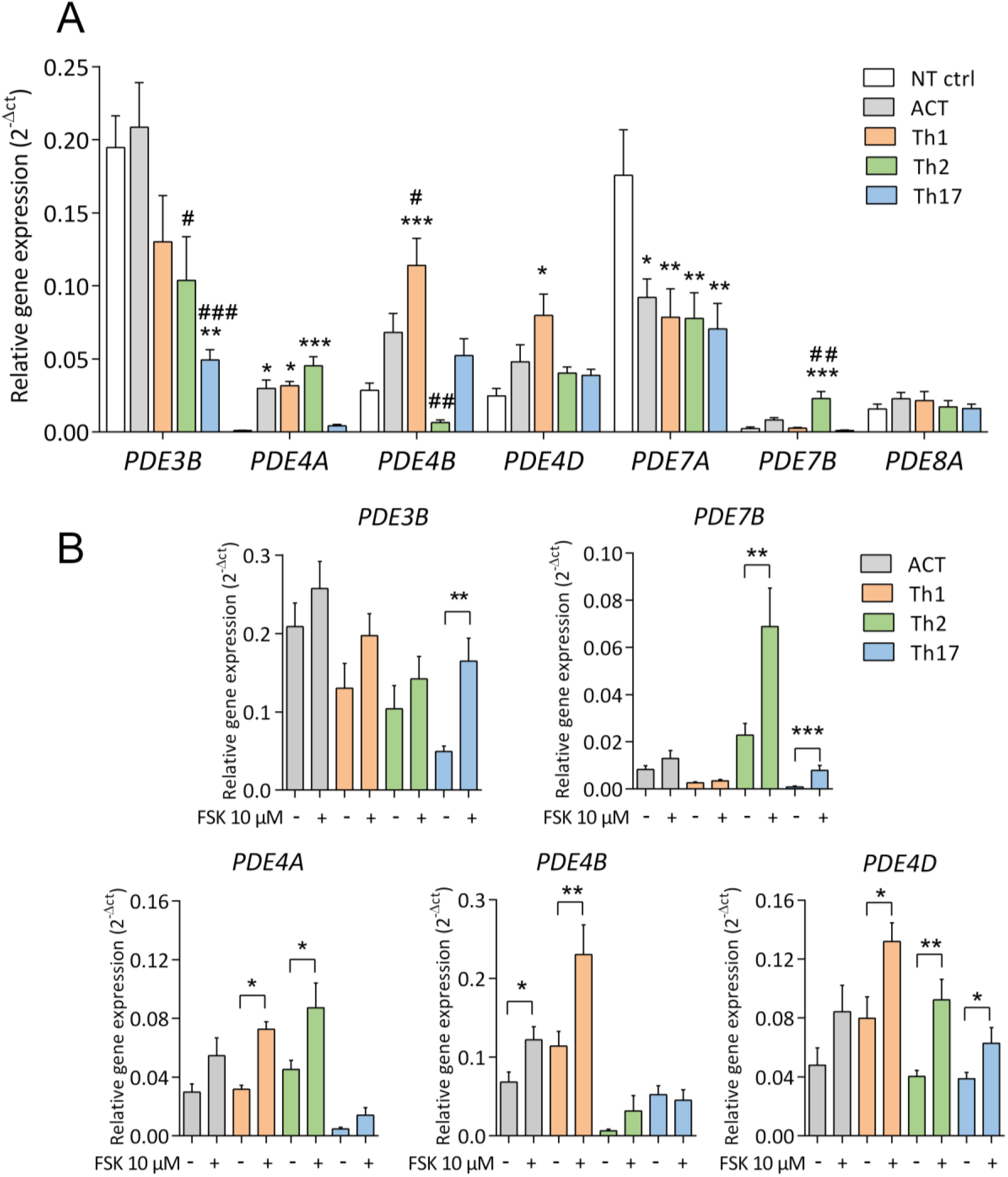
Analysis of phosphodiesterase (PDE) isoform expression in Th1, Th2 and Th17 cells. **A**. Expressions of detectable PDE isoforms (PDE3B, PDE4A, PDE4B, PDE4D, PDE7A, PDE7B and PDE8A) in the Th subsets (Th1, Th2 and Th17) and both non-treated (NT) and activated (ACT) controls. Statistical significance was examined by one-way ANOVA (* vs NT ctrl, # vs ACT) **B**. Analysis of FSK-mediated changes in the expressions of selected PDE isoforms. Statistical significance was examined by unpaired two-tailed Student’s t-test. All data are shown as mean ± SEM, n=4-9, *p < 0.05, **p < 0.01, ***p < 0.001

FSK mediated an increase in *PDE3B* expression independently of the type of Th subset; however, this increase was significant only in Th17 cells (**Figure 3B**). On the other hand, cells that differentiated into all three subsets showed significant elevations of *PDE4D*. The expression of *PDE4B* was significantly elevated in ACT and Th1. The expression of *PDE4A* was significantly elevated in the Th1 and Th2 subsets. *PDE7B* showed a significant increase in the Th2 and Th17 subsets. There were no significant changes in the expressions of *PDE7A* or *PDE8A* after FSK stimulation (**Supplementary data 5B**).

### Effect of FSK-mediated AC activation on Th cell differentiation

To describe the complexity of the FSK-mediated modulation of Th1, Th2, and Th17 cell differentiation, data from scRNAseq analysis were combined with analyses of the expressions of specific genes and the production of specific cytokines. Firstly, employing the scRNAseq analysis, the UMAP algorithm (uniform manifold approximation and projection for dimension reduction) with a resolution of 0.9 was applied to achieve a compromise between the bordered clusters and cell distribution, this value for the resolution being in the optimal range suggested by Seurat (RStudio package for quality control). The dataset was divided into 12 specific clusters (**Figure 4A**). Interestingly, some of these clusters corresponded to cells that differentiated into the Th1, Th2, and Th17 subsets, or Th1, Th2, and Th17 cells treated with 10 μM FSK (marked as Th1+FSK, Th2+FSK, Th17+FSK) that were labelled by cell surface feature barcodes (**Figure 4B**) proving differentiation into the intended Th subsets. Apparently, cluster 0 belonged to the Th1 and Th1+FSK subsets, cluster 1 to the ACT and ACT+FSK subsets, cluster 2 to NT ctrl, cluster 3 to the Th2 subset, and cluster 4 to the Th2+FSK subsets, while cluster 5 corresponded mostly to the Th17 subset and clusters 8 and 9 were formed by the Th17+FSK subsets, as described later. In addition, some of the clusters (6, 7, 10, and 11) were formed independently of the intended Th cell differentiation marked by sample labelling. Clusters 6 and 10 were considered to be cells in transient differentiation stages that do not possess enough markers to allow them to be included into the Th1 or Th17 clusters. Interestingly, a small group of *FOXP3*+ cells with the transcriptional profile of Treg cells could be identified, creating cluster 11 (**Figure 4A, Supplementary data 6B**). This cluster comprised cells from various Th differentiation lines, with the highest percentage of Th1 and FSK-treated Th1 cells (45.5 %)(**Supplementary data 6C**). At the same time, this cell population comprised only a small fraction of Th1 cells (6 %) and FSK-treated Th1 cells (9.5 %).

**Figure 4.**
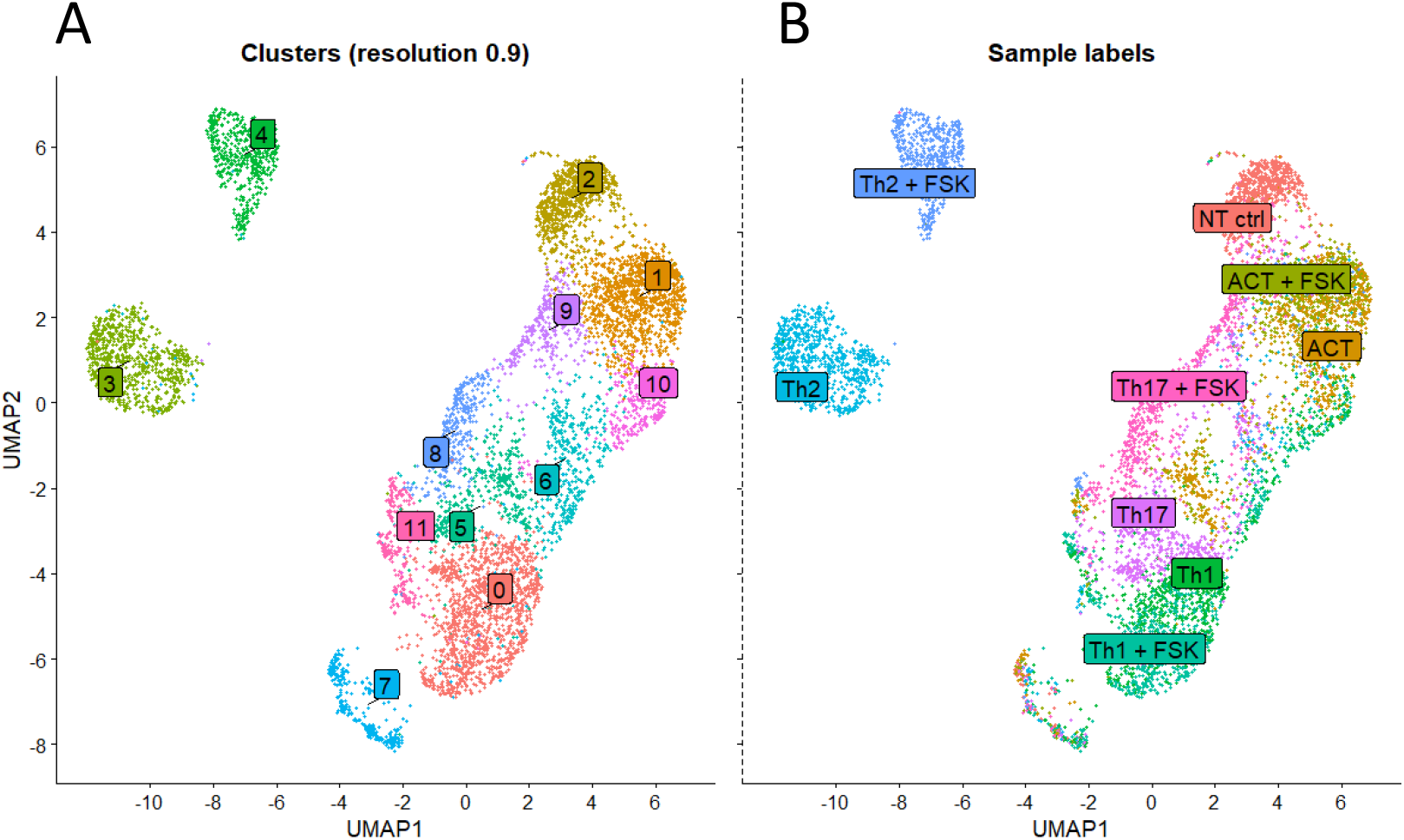
UMAP visualization of clusters. **A**. Automatic clustering based on the UMAP algorithm. Colors represent cells in the 12 clusters defined using top variable genes in the unsupervised clustering. Labels represent the numbers assigned to the clusters distinguished by the FindNeighbours function (Seurat) under a resolution of 0.9; **B**. Visualisation of sample-specific labeling (hashtag antibodies). Colors and labels represent cells in clusters corresponding to the particular cell feature barcodes that were used for multiplexing during sample preparation.

The effect of FSK-mediated AC activation on Th1, Th2, and Th17 subset differentiation is described in detail in the following sections.

#### FSK-induced elevation of Th1-specific markers reinforcing the Th1 cell phenotype

The effects of FSK on Th1 differentiation were primarily examined on the basis of the effects on the gene expressions of *TBX21* and *ANXA3* and on the production of the effector cytokine IFN-γ, typical for the Th1 subset. FSK significantly increased the expression of *ANXA3* as well as the accumulation of IFN-γ in a dose-dependent manner (**Figure 5B and C**). However, *TBX21* expression was only minimally elevated (**Figure 5A**). The detailed scRNASeq analysis of extracted Th1 datasets showed no apparent differences between FSK-treated (10 μM) and non-treated differentiated Th1 cells, according to their localization in the UMAP (**Figure 5D**). However, HeatMap visualization of the 15 most differentially expressed genes induced by FSK showed some upregulation in the expressions of the genes encoding the following – tumor necrosis factor-activated receptor (*TNFRSF18*), metal ion binding protein (*MT2A*), MAP kinase phosphatases (*FURIN, DUSP4*), and phosphodiesterase *PDE4B*. Moreover, the gene for granzyme B (*GZMB*) appeared among the most strongly upregulated genes induced by FSK (**Figure 5E**).

**Figure 5.**
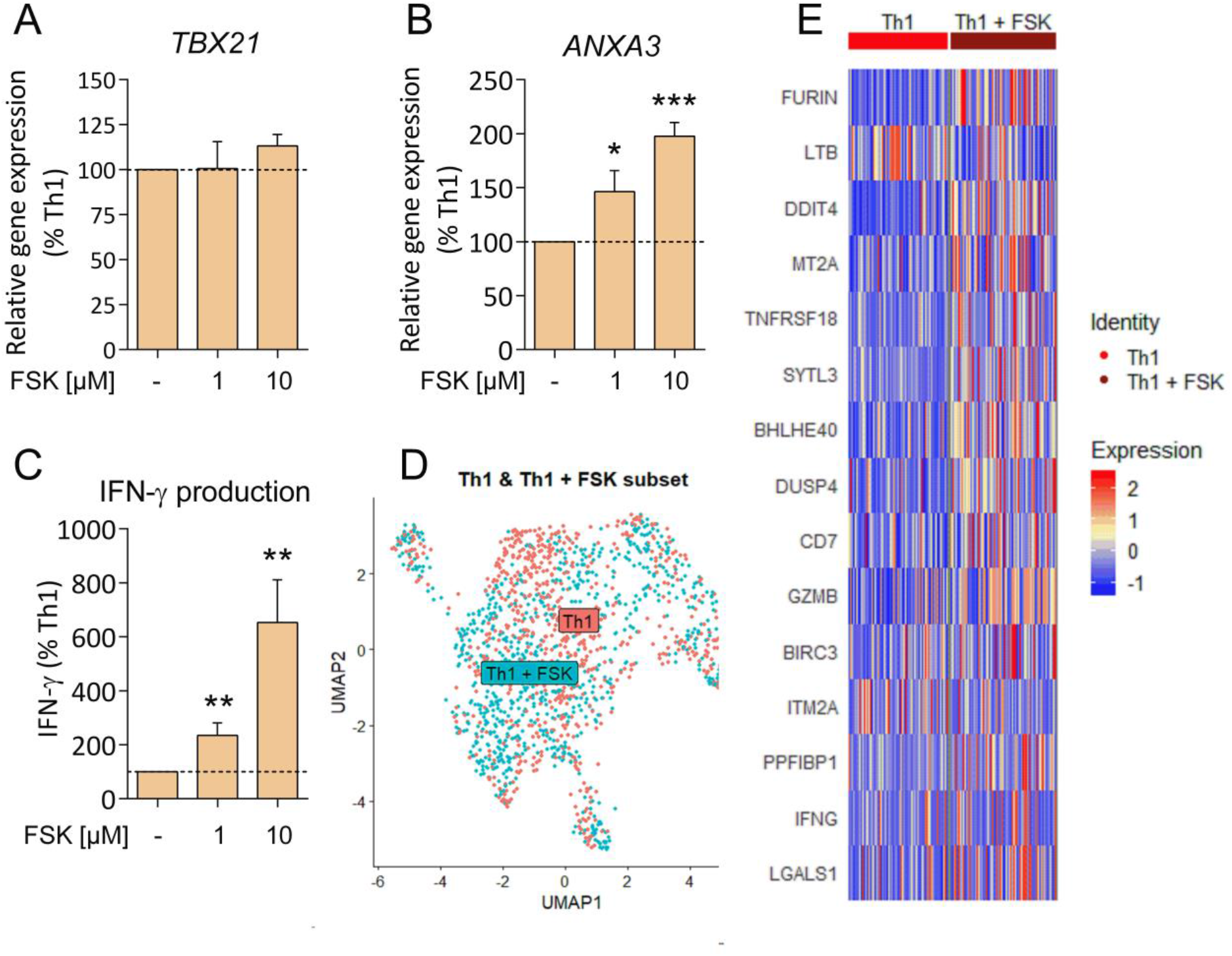
FSK promotes Th1 differentiation via the induction of Th1-specific markers. **A-C**. Effect of 1 or 10 μM FSK on the expressions of two selected Th1-specific genes (TBX21 and ANXA3) and production of the effector cytokine IFN-γ. Data are expressed as percentage of Th1 cells. Statistical significance was examined by one-sample t-test, separately comparing the effect of each experimental treatment with Th1 cells. Mean ± SEM, n=5-9, *p < 0.05, **p <0.01,***p < 0.001. **D**. scRNAseq UMAP visualization of Th1 subset (red dots) and Th1 cells upon treatment with 10 μM FSK (Th1+FSK subset, blue dots) **E**. HeatMap visualization of the 15 most differentially expressed genes in the Th1 (first column) and Th1+FSK subsets (second column). Different colors correspond to the scaled expression (Z-score).

#### Th2 cell differentiation was strongly supported by FSK

Similarly to Th1, FSK induced upregulation in the gene expressions of specific transcription factor *GATA3* and *LIMA1* as well as the production of effector cytokine IL-13 (**Fig 6A-C**). The detailed scRNAseq analysis of extracted Th2 datasets resulted in a clear division of samples into corresponding clusters with only small overlaps (**Figure 6D**). FSK (10 μM) treatment led to the upregulation of genes, encoding e.g. the heat-shock protein (*HSPA8*) and chemokine ligand (*XCL1*). The most significantly downregulated genes were *WARS, GARS, and SAR*S, encoding aminoacyl-tRNA ligases, and genes *SLC3A2, PSAT1, and PHGDH*, encoding proteins involved in amino-acid metabolism (**Figure 6E**).

**Figure 6.**
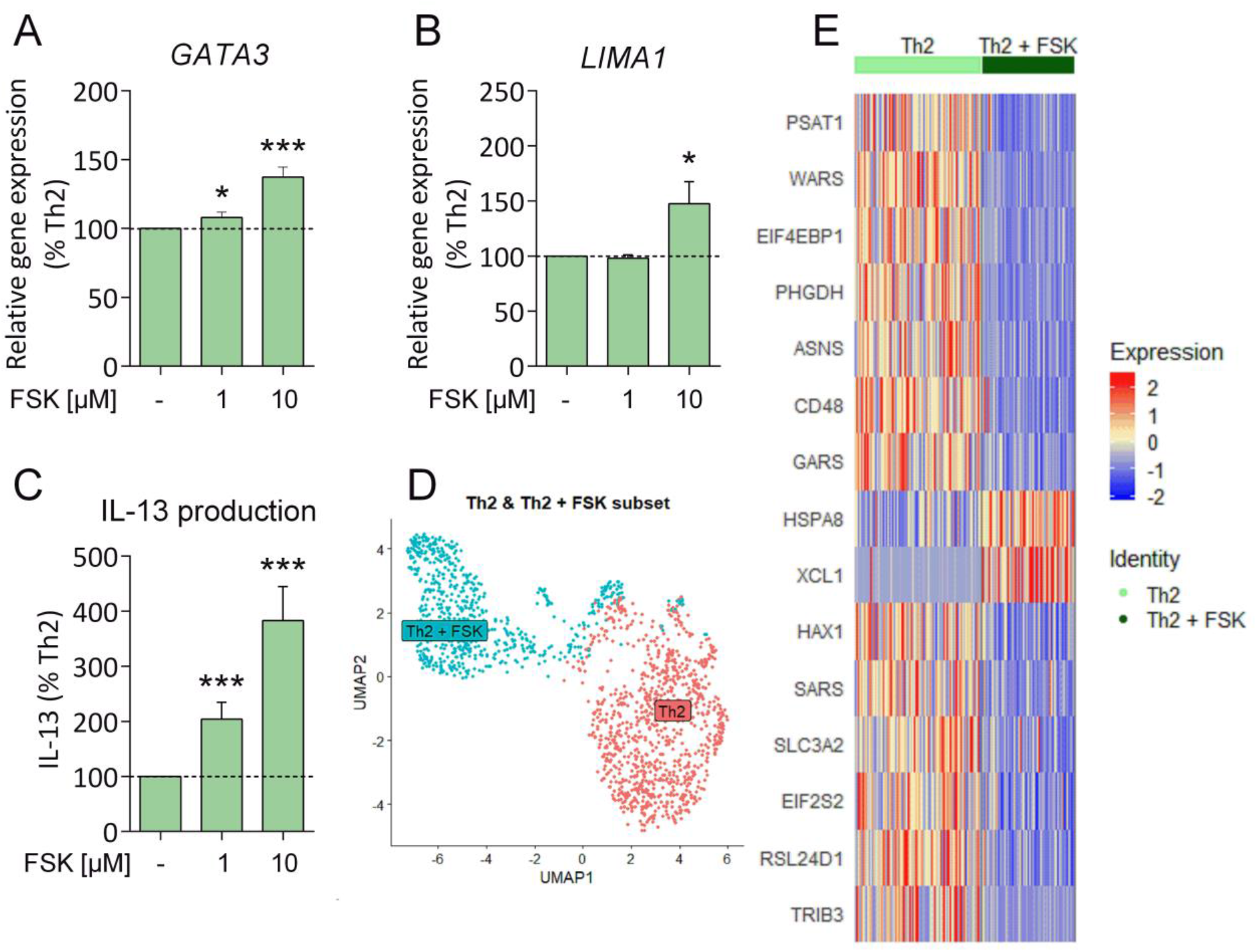
FSK supports Th2 differentiation. **A-C**. Effect of 1 or 10 μM FSK on the expressions of two selected Th2-specific genes (GATA3 and LIMA1) and production of the effector cytokine IL-13. Data are expressed as percentage of Th2 cells. Statistical significance was examined by one-sample t-test, separately comparing the effect of each experimental treatment with Th2 cells. Mean ± SEM, n=5-8, *p < 0,05, **p < 0,01, ***p < 0,001. scRNAseq UMAP visualization of Th2 subset (red dots) and Th2 cells upon treatment with 10 μM FSK (Th2+FSK subset, blue dots) **E**. HeatMap visualization of the 15 most differentially expressed genes in the Th2 (first column) and Th2+FSK subsets (second column). Different colors correspond to the scaled expression (Z-score).

Moreover, scRNAseq analysis pointed out two genes whose expressions were dramatically altered upon FSK treatment (**Figure 7A and B**). PPARγ, encoded by the *PPARG* gene, is the regulator of fatty acid metabolism, which plays an important role in Th2 differentiation. The data showed that FSK induced significant elevation of the gene **(Supplementary data 7C)** as well as the protein expression of PPARγ (**Figure 7C**). This PPARγ elevation correlated with the repression of IL-2 production (**Supplementary data 7B**) and downregulation of transcription factor NFAT (**Supplementary data 7A**). Further, the expression of Sestrin2 (*SESN2* gene), the upstream regulator of mTOR, displayed significant downregulation upon FSK treatment (**Figure 7D**), suggesting the importance of the mTOR/Akt pathway in Th2 differentiation. In addition, the data from scRNAseq revealed the impaired expression of other mTOR/Akt-related genes, e.g. *PDK1* (**Supplementary data 7A**).

**Figure 7.**
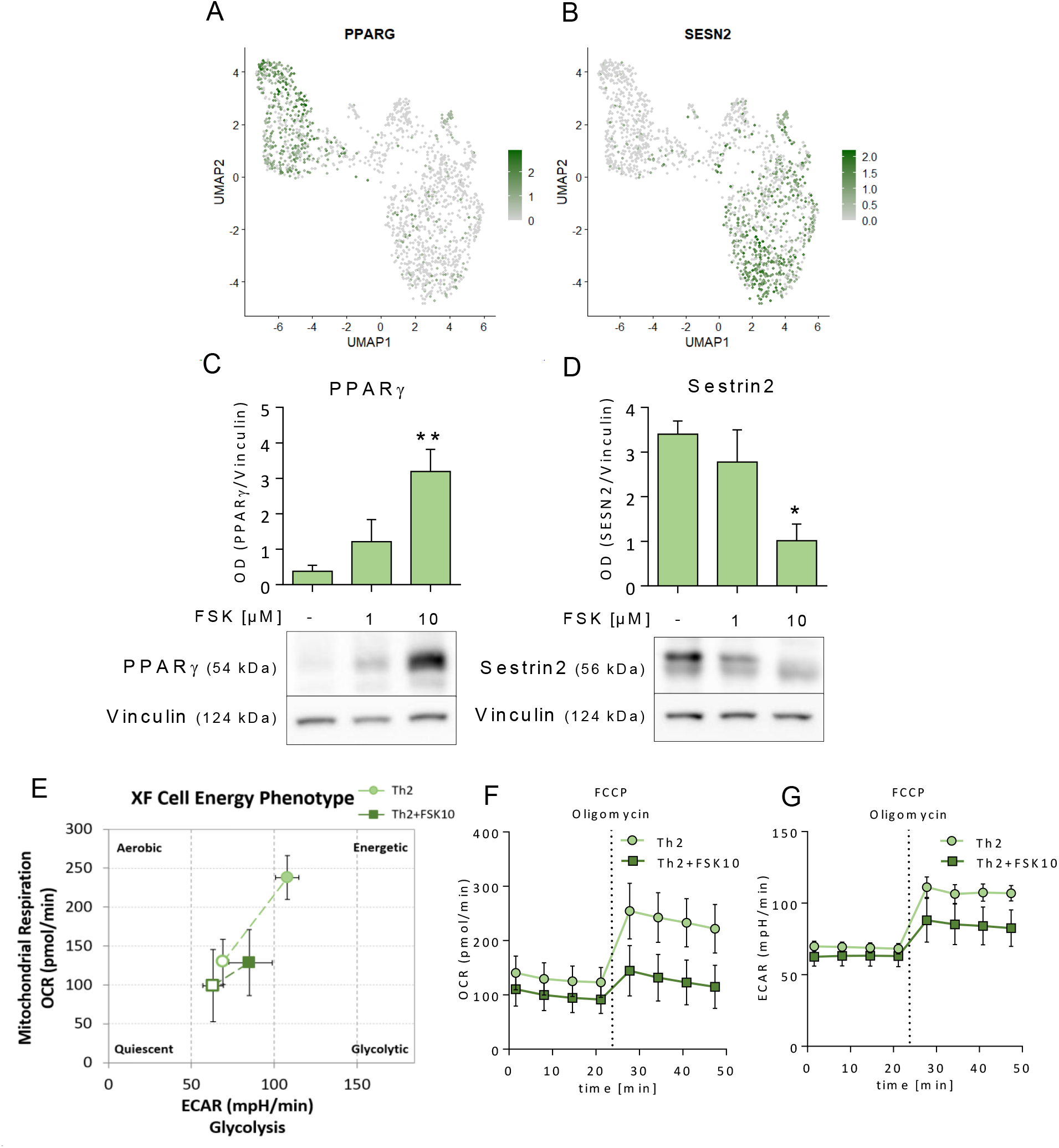
Analysis of FSK influence on Th2 metabolism. **A,B**. scRNAseq UMAP visualization of SESN2 and PPARG positive cells (green) in the extracted Th2 and Th2+FSK subsets. Dark and light green colors correspond to the scaled expression (Z-score); **C,D**. Analysis of protein expressions of Sestrin2 and PPARγ in Th2 cells and 1 or 10 μM FSK-treated Th2 cells. Statistical significance was examined by one-way ANOVA. Mean ± SEM, n=5, **p < 0.01; **E-G**. Analysis of mitochondrial respiration (OCR, oxygen consumption rate) and glycolysis (ECAR, extracellular acidification rate) measured using the Seahorse XF Cell Energy Phenotype test. The differences between baseline activity (open symbols) and stressed activity (closed symbols), which was induced by a simultaneous injection of the stressor compounds FCCP and oligomycin (both 1 μM), is shown. Mean ± SEM, n=4.

The overall effect of FSK on Th2 cell metabolism during the differentiation process was further analyzed using Agilent Seahorse XF technology, which allows simultaneous measurement of the two major energy-producing pathways - mitochondrial respiration (OCR, oxygen consumption rate) and glycolysis (ECAR, extracellular acidification rate). In contrast to differentiated Th2 cells, the cells treated with FSK (10 μM) revealed a metabolic switch, as could be seen in stressed conditions induced by FCCP and oligomycin (**Figure 7E**). The exposure of Th2 cells to FSK did not change baseline activity but caused a decrease in the utilization of both pathways under metabolic stress - severely in the case of OCR but only mild in the case of ECAR (**Figure 7F, Supplementary data 7D,E)**. These data indicate suppression of the oxygen-dependent metabolic potential of Th2 cells upon FSK treatment (**Figure 7G, Supplementary data 7F)**. These changes suggest that FSK inhibited proliferation. This conclusion was supported by a proliferation assay showing the impairment of Th2 proliferation upon FSK treatment (**Supplementary data 1B**.).

Due to the complexity of the results regarding the FSK-mediated modulation of Th2 differentiation, we decided to elucidate the mechanism further. The P-site inhibitor 2’,5’-dideoxyadenosine (ddAdo) was used to block the activity of ACs, allowing confirmation of the hypothesis that FSK-mediated AC activation is responsible for the aforementioned phenomena. The effect of the AC modulators FSK and ddAdo on the gene expressions of subset-specific markers *GATA3* and *LIMA1*, and selected metabolism-related genes *PPARG* and *SESN2* was studied in differentiated Th2 cells. ddAdo (100 μM) was added to Th cell cultures during the differentiation process 20 h after activation. As expected, the inhibition of ACs mediated by 100 μM ddAdo 30 min prior to FSK (10 μM) stimulation resulted in the suppression of cAMP production (**Supplementary data 8**.). Interestingly, the supportive effect of FSK on the gene expressions of the Th2-specific markers *GATA3* and *LIMA1* was suppressed when ACs were firstly inhibited by ddAdo (**Figure 8A and B**). The gene expression of *GATA3* was mildly decreased (**Figure 8A**), while the downregulation of *LIMA1* expression was significant (**Figure 8B**). Th2 cells treated with FSK and ddAdo revealed a significantly lower gene expression on the part of *PPARG* compared to cells treated with the FSK only (**Figure 8C**). The downregulation of *SESN2* expression upon FSK treatment was also disrupted using the inhibitor ddAdo, this resulting in an increase in the gene expression (**Figure 8D**).

**Figure 8.**
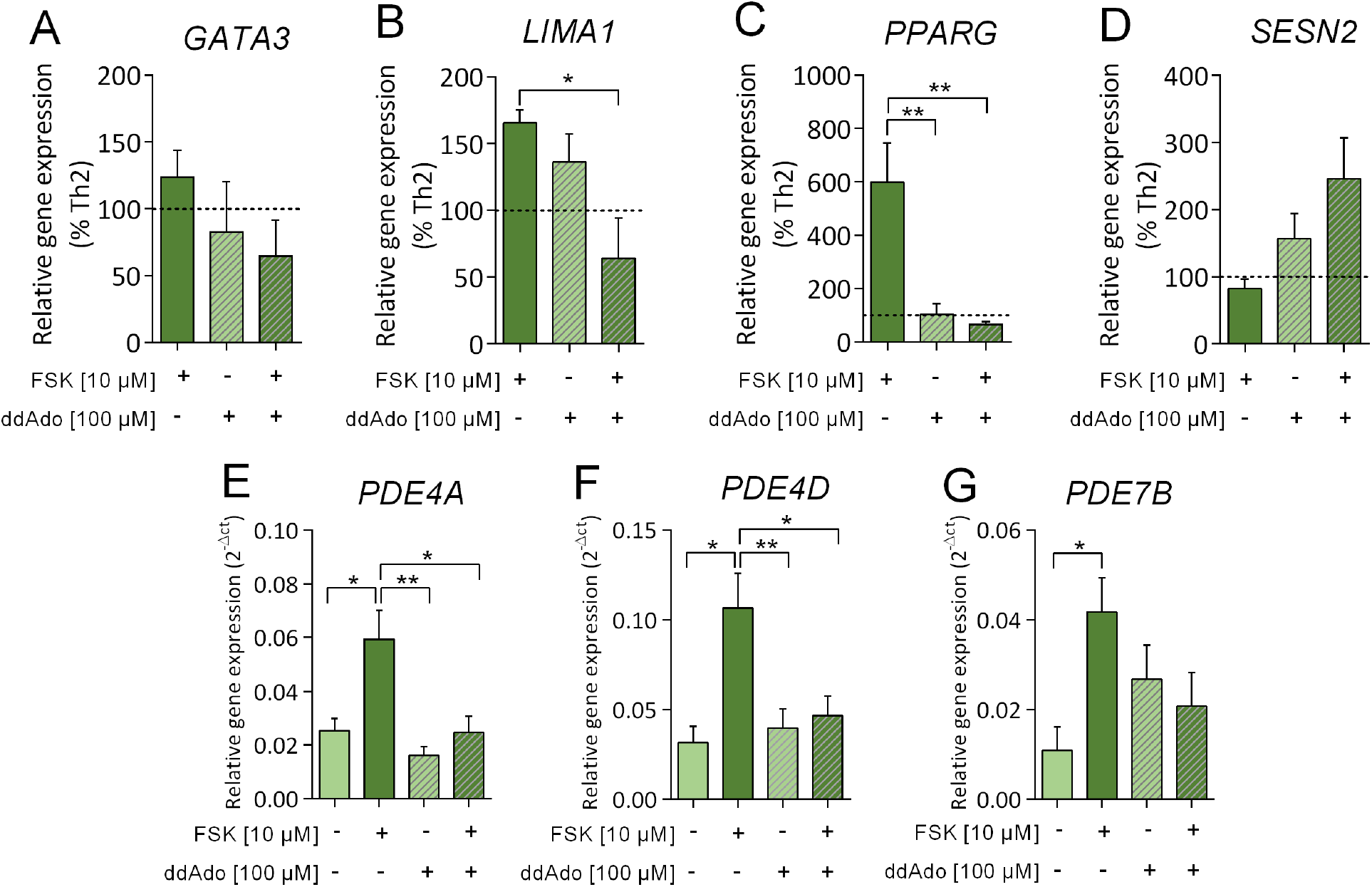
Effect of ddAdo-mediated inhibition of AC on Th2 differentiation. **A-D**. Effect of 10 μM FSK and 100 μM ddAdo on the expression of two selected Th2-specific genes (GATA3 and LIMA1) and the studied genes PPARG and SESN2. Dotted line represents 100% (Th2 cells). Data are expressed as percentage of Th2 cells. **E-G**. Analysis of gene expression of selected PDE isoforms (PDE4A, PDE4D and PDE7B) in differentiated Th2 cells upon FSK (10 μM) and ddAdo (100 μM) treatment. Statistical significance was examined by one-way ANOVA. Mean ± SEM, n=3-5. * p < 0.05, ** p < 0.01

In addition, we focused on the analysis of *PDE4A*, *PDE4D* and *PDE7B* gene expression, which were shown to be significantly elevated in Th2 cells upon FSK treatment and can be suggested to have an important role in this process (**Figure 8E-G**). The gene expressions of these isoforms in Th2 cells treated with ddAdo or a combination of ddAdo and FSK did not differ from those in non-treated Th2 cells. Interestingly, unlike in Th2 cells treated with FSK alone, the effect of FSK on the gene expressions of *PDE4A, PDE4D, and PDE7B* was disrupted by the inhibitor ddAdo (**Figure 8E-G**). Overall, Th2 cells treated with ddAdo or ddAdo and FSK did not achieve the same levels of PDEs as those treated with FSK only. However, the difference was significant only in the case of *PDE4A* and *PDE4D*.

#### FSK induced the suppression of Th17 differentiation, leading to the formation of a specific CXCL13+ population

Interestingly, FSK mediated downregulation of the expression of the Th17-specific markers *RORC2* and *PALLD*, as well as production of the effector cytokine IL-17A (**Figure 9A-C**). Further detailed scRNAseq analysis of the extracted Th17 dataset showed a division into two clusters with partial overlap (**Figure 9D**). HeatMap visualization of the 15 most differentially expressed genes (**Figure 9E**) showed that only very few genes were upregulated after FSK treatment (10 μM) and that most of the genes underwent significant downregulation. Importantly, the expression of the Th17-specific signature gene *PALLD* underwent FSK-mediated downregulation, which was confirmed by data obtained via RT-qPCR using various donors (**Figure 9B**). Moreover, FSK treatment during the Th17 differentiation process resulted in a reduction in proliferation (**Supplementary data 1B**.). In cluster 8 of UMAP (**Figure 4**), *CXCL13* was found to be one of the most upregulated genes after FSK exposure. These cells corresponded to the sample Th17+ FSK (**Supplementary data 6A**.). Further analysis by RT-qPCR showed a significant increase in *CXCL13* gene expression in Th17 cells upon FSK treatment (**Figure 9F**), which was confirmed at the protein level (**Figure 9H**).

**Figure 9.**
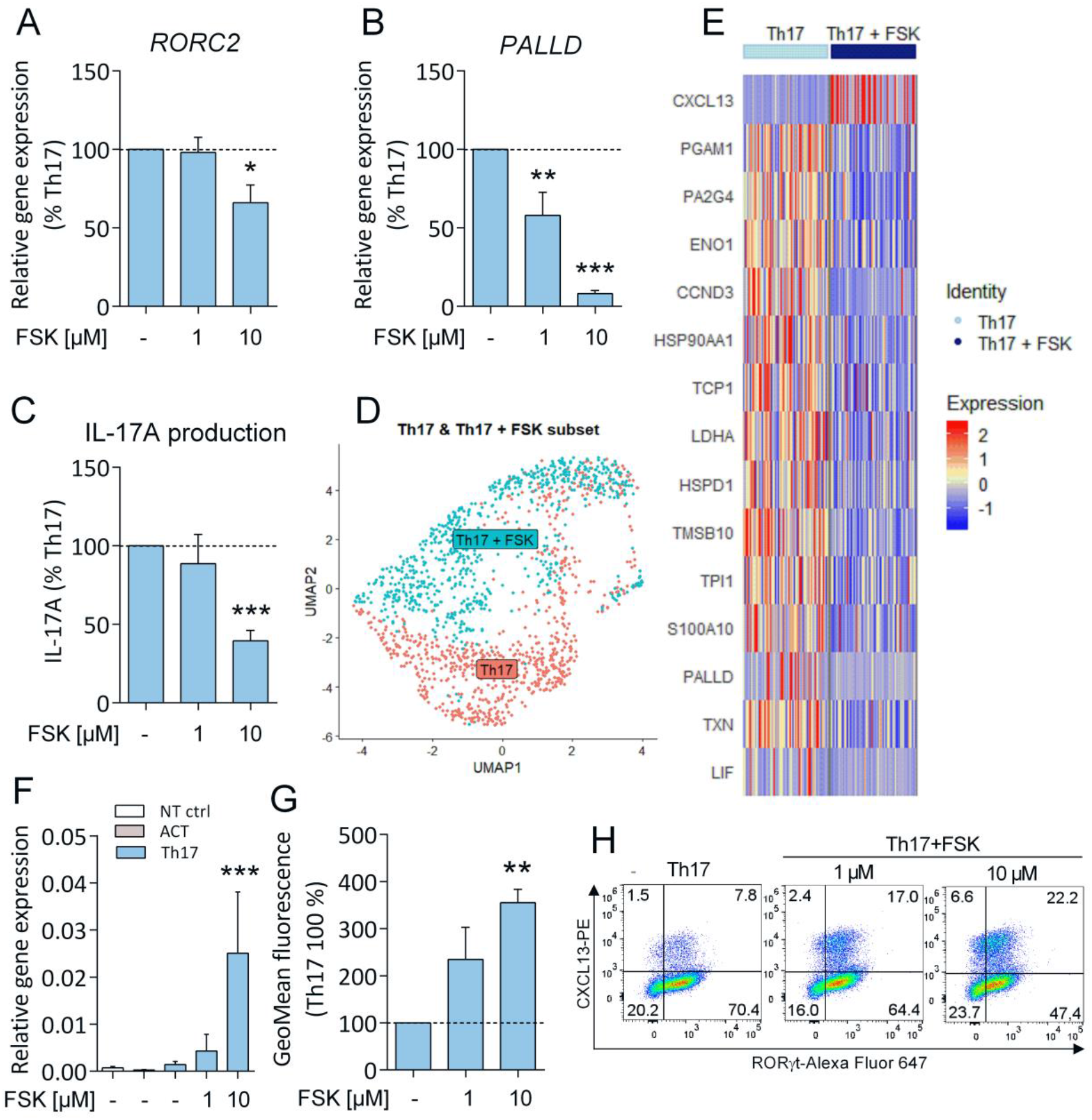
Forskolin-mediated impairment of Th17 differentiation led to specific CXCL13+ cell formation. **A-C**. Effect of 1 or 10 μM FSK on the expression of two selected Th17-specific genes (RORC2 and PALLD) and production of the effector cytokine IL-17A. Data are expressed as percentage of Th17 cells. Statistical significance was examined by unpaired two-tailed Student’s t-test, separately comparing the effect of each experimental treatment with Th17 cells. Mean ± SEM, n=5-8, *p < 0.05,**p < 0,01, ***p < 0.001. **D**. scRNAseq UMAP visualization of Th17 subset (red dots) and Th17 cells upon treatment with 10 μM FSK (Th17+FSK subset, blue dots). **E**. HeatMap visualization of the 15 most differentially expressed genes in the Th17 (first column) and Th17+FSK subsets (second column). Different colors correspond to the scaled expression (Z-score). **F**. Analysis of relative gene expression of CXCL13 in Th17 cells and Th17 cells treated with 1 or 10 μM FSK. Mean ± SEM, n=6, ***p < 0.001. **G**. Flow cytometry analysis of CXCL13 in Th17 cells and Th17 cells treated with 1 or 10 μM FSK. Geometric Mean of fluorescence is expressed as a percentage of Th17 cells. Mean ± SEM, n=4, ** p < 0.01. **H**. Representative dot plot visualizing CXCL13 and RORγt expression. Numbers in plots indicate the percentage of cells in each area.

## 3. Discussion

This study shows the significance of the FSK-mediated reinforcement of the differentiating Th1 cell phenotype, the modulation of the metabolic phenotype in Th2 cells, and the suppression of Th17 differentiation. The expression of some of the AC isoforms as well PDEs were changed in differentiating Th cells – however, without any uniform pattern. The data clearly show differences in the influence of FSK on the differentiation of distinct Th subsets.

The effect of FSK was analyzed using the *in vitro* differentiation of human naïve Th cells into Th1, Th2, and Th17 subsets, which was verified via analyses of transcription factor expressions and the production of the main effector cytokines specific to distinct subsets. The increase in Th cell proliferation confirmed the functional activation of cells by selected stimuli (**Supplementary data 3**) (***Luckheeram et al., 2012***). As expected, T-bet (*TBX21*) and IFN-γ were found to be specific to Th1 cells, GATA3 (*GATA3*) and IL-13 to Th2 cells, and RORγt (*RORC2*) and IL-17A to Th17 cells, according to analyses performed on multiple donors. In addition, scRNAseq analysis provided detailed transcriptomic data on the differentiated Th cell subsets, allowing the selection of the signature genes *ANXA3* and *RUNX3* for the Th1 subset, *LIMA1* and *MAOA* for the Th2 subset, and *PALLD* and *IL23R* for the Th17 subset on the basis of recent studies ***(Cano-Gamez et al., 2020, Tibbitt et al., 2019)***. In combination with the classic approach, the novel approach of scRNAseq clarified the targeted differentiation of Th cells.

To gain a perspective on the expression of the two key enzyme families regulating intracellular cAMP levels, ACs and PDEs were evaluated. As expected, expression of the three main immune cell-specific AC isoforms, *ADCY3*, *ADCY7* and *ADCY9* (***Dessauer et al., 2017***), was detected in Th cells. Moreover, *ADCY1* also appeared among detectable isoforms upon Th cell activation. Surprisingly, an observable shift in the ratio of *ADCY3* and *ADCY7* expression was found upon Th cell activation. Whereas *ADCY7* was the dominant AC isoform expressed in Th naïve cells, *ADCY3* appeared to be dominant in activated Th cells. However, recent studies declared *ADCY7* to be essential for the functioning of immune cells in the host organism, as was observed in an *ADCY7* knockout murine model (***Duan et al., 2010***).

The direct opposite of ACs are PDE enzymes that are responsible for cAMP and cGMP degradation. PDE families in the immune system comprise PDE3, 4, 7 and 8 (as reviewed in ***(Raker et al., 2016, Wehbi and Tasken, 2016***)). The expression of *PDE7A* was specifically assigned to Th naïve cells (***Nueda et al., 2006***), which is consistent with our data. *PDE4B* was primarily expressed in Th1 cells, while *PDE4A* and *PDE7B* were specific to Th2 cells. Whereas *PDE4B* and *PDE7B* maintained their specificity to the Th1 and Th2 subsets, even upon FSK treatment, *PDE4A* appeared to be upregulated upon FSK treatment in Th1 and Th2, except for Th17. It can be speculated that the upregulation of PDE isoforms serves as a confirmation of the successful activation of the cAMP signaling pathway, since the downstream molecule PKA was shown to be associated with upregulated PDE as a part of a negative feedback loop, as mentioned in the case of PDE4D ***(Liu et al., 2000***). Overall, PDE expression in differentiating Th cells revealed high variability among the various Th subsets, which did not allow firm conclusions to be drawn about the role of particular PDEs.

Regarding the effect of FSK on the studied Th1 cell subset, the obtained data showed an FSK-mediated reinforcement of Th1 differentiation. The overlap between the Th1 subset and Th1 cells differentiated in the presence of FSK, documented by scRNAseq analysis, suggested that the typical phenotype of the Th1 subset was not altered upon FSK treatment. Furthermore, elevation in the expression of the Th1 specific signature genes *ANXA3* and *RUNX3* was observed, as well as an increase in IFN-γ production. However, the expression of *TBX21* displayed only limited upregulation upon FSK treatment. These results are in accord with a recent study showing the activation of the cAMP signalling pathway stimulated by the direct binding of PGE2 (***Yao et al., 2013***) or PGI2 (***Nakajima et al., 2010***) to GPCR, which led to the promotion of Th1 differentiation and effector functions. Interestingly, enrichment analysis using the Gene Ontology Molecular Function Database 2018 (GO-MF, (***Mi et al., 2019***)) revealed increased expression in genes participating in metal ion binding (*MT2A*, *MCTP2*, *SYTL3*), MAP kinase phosphatase activity (*FURIN*, *DUSP4*), and phosphodiesterase activity (*PDE4B*). Moreover, the gene for granzyme B (*GZMB*) appeared among the genes most strongly upregulated by FSK treatment, which is consistent with the effector functions of the Th1 subset (***Kreutzman et al., 2014***). Finally, enrichment linked the FSK-mediated cAMP increase with the elevated expression of the *TNFRSF18* gene for the TNFα receptor. Interestingly, this gene was previously connected to the transcription profile of the Treg subset and assigned to Treg-specific signature genes ***(Mahmud et al., 2014, Petrillo et al., 2015)***.

Regarding the effects of the FSK-mediated elevation of cAMP on Th2 cell differentiation, both scRNAseq and RT-qPCR showed the up-regulated expression of the gene encoding the main Th2-specific transcription factor *GATA3*, as well as other Th2-specific signature genes, namely *LIMA1* and *MAOA*, suggesting support for the idea of Th2 differentiation occurring by means of FSK treatment. This was further supported by the enhanced production of the Th2 effector cytokine IL-13 by FSK. These findings are in agreement with the study of Elliott et. al presenting a support of Th2 differentiation by elevated cAMP ***(Elliott et al., 2001***). It is known that the Th2 cell subset employs a strong differentiation machinery that enhances itself through an efficient positive feedback loop (***Zhu, 2010***). Interestingly, it was previously shown that an elevated cAMP level led to the upregulation of IL-4 production and therefore accelerated Th2 differentiation mechanisms (***Tokoyoda et al., 2004***). A similar effect could also be suggested here, showing FSK-mediated cAMP to enhance Th2 differentiation. The increase in Th2 specific markers in the presence of FSK was accompanied by a decrease in the proliferation of FSK-treated Th2 differentiating cells. This further supports the higher differentiation status of Th2 cells differentiated in the presence of FSK.

Here, the impact of FSK was so remarkable that it separated the differentiating Th2 cells into two distinct clusters with minimal overlap, as shown by the scRNAseq analysis. A decrease in the gene expressions of enzymes involved in cell metabolism was the most crucial factor in the separation of the FSK-treated Th2 cluster, according to the enrichment analysis of the differentially expressed genes using the GO-MF. Among the most significantly impaired cellular processes were aminoacyl-tRNA ligase activity (*WARS, GARS, SARS*) and amino-acid transport (*SLC3A2*). Also, the decreased expressions of *PSAT1* (phosphoserine aminotransferase 1) and *PHGDH* (phosphoglycerate dehydrogenase) indicated a decrease in serine metabolism. Interestingly, Jun et al. reported that the expression of the *PHGDH* gene, dictated by IL-2R signaling, is crucial for DNA synthesis during the S phase of activated T cells (***Jun et al., 2014***), while its down regulation in FSK-treated cells correlates with the observed suppression of the proliferation of these cells.

The differentiation of Th cells is connected with a change in cell metabolism towards glycolysis ***(Stark et al., 2019***). A decrease in the spare respiration capacity could promote the glycolytic metabolism in FSK-treated Th2 cells. The high rate of glycolysis in differentiating T cells requires activation of the kinase mTOR (***Stark et al., 2019***). Interestingly, we observed downregulation of the expression and protein levels of Sestrin2, a negative regulator of mTOR (***Ben-Sahra et al., 2013***), upon FSK-mediated cAMP elevation. We can speculate that this could contribute to a possible mechanism of FSK-mediated change in cell metabolism. Interestingly, mTOR activity is also essential for the upregulation of lipid metabolism, especially the lipid synthesis pathway, which is an important aspect of Th cell activation (***Stark et al., 2019***). The importance of FSK-mediated cAMP in the modulation of lipid metabolism was documented by a highly significant increase in the expression as well as protein levels of PPARγ, the key regulator of lipid metabolism in cells, upon FSK treatment. PPARγ becomes highly expressed specifically in Th2 cells and likely regulates fatty acid metabolism later in the Th2 cell differentiation program (***Stark et al., 2019***). However, PPARγ is described as a rather controversial player with opposing effects on distinct Th2-specific effector functions. According to the literature, PPARγ is involved in the suppression of IL-2 production, which is orchestrated by the NFAT transcription factor (***Hernandez-Quiles et al., 2021***). Interestingly, this correlation was confirmed in our study, with Th2 cells differentiated in the presence of FSK revealing a lower IL-2 production.

The causality between FSK-mediated cAMP production and the aforementioned effect of FSK on Th2 differentiation was demonstrated by employing ddAdo, an AC inhibitor reported to reduce FSK-mediated cAMP production in T cells (***Schillace et al., 2007***). The effect of FSK on the gene expressions of Th2 subset-specific markers and selected metabolism-related genes was disrupted by ddAdo treatment. Unlike in the case of Th2 cells treated with FSK only, the gene expressions of *GATA3*, *LIMA1*, and *PPARG* in cells treated by ddAdo were suppressed, while *SESN2* gene expression was elevated. Moreover, ddAdo suppressed the FSK-mediated elevation of *PDE4A*, *PDE4D* and *PDE7B* expression in Th2 cells. Thus, the gene expressions of *PDE4A*, *PDE4D* and *PDE7B* remained at the same levels as in non-treated Th2 cells. This observation suggests the importance of *PDE4A*, *PDE4D* and *PDE7B* in the cAMP-mediated regulation of Th2 differentiation.

In contrast to the situation with the Th1 and Th2 subsets, FSK mediated the suppression of all selected markers of differentiating Th17 cells. Decreases in the expressions of the transcription factor RORγt (*RORC2*) and the selected signature genes *PALLD* and *IL23R*, and the inhibition of IL-17A production were observed. Moreover, an analysis of the enrichment of differentially expressed genes, based on the GO-MF, showed the downregulation of genes encoding heat-shock proteins (*HSP90AA1, HSPD1*), proliferation-associated molecules (*PA2G4, CCND3*), and proteins connected to calcium signaling (*S100A10*). This suggests the impairment of Th17 cell differentiation at different levels of cell regulation. Surprisingly, these findings suggesting the repressive effect of FSK-mediated cAMP elevation on Th17 cell differentiation appear to contrast with other studies, though these other studies employed various types of PGs (PGE2, PGI2) to trigger the cAMP signalling pathway in earlier stages of cell differentiation ***(Boniface et al., 2009, Liu et al., 2013, Sakata et al., 2010***). In contrast, our study was based on the highly specific and direct activation of AC by FSK, which stimulates the cAMP signalling pathway at a more downstream level. Interestingly, our results appear to be consistent with the conclusions of studies focused on the effects of cAMP elevation achieved by PDE inhibition. The inhibitors of PDE4, apremilast (***Schafer et al., 2014***) or Zl-n-91 (***Ma et al., 2008***), significantly decreased the production of Th17-specific effector cytokine, IL-17A. Interestingly, the differential expression analysis pointed to one specific gene, *CXCL13*, which was strongly upregulated upon FSK treatment of Th17 cells, an increase also confirmed at the protein level. CXCL13 provides important costimulatory signals to B cells that should be produced mainly by stromal cells and follicular dendritic cells (***Kazanietz et al., 2019***). Interestingly, Takagi et al. reported that specific subclones of Th17 cells, but not Th1 or Th2 cells, can also express CXCL13 (***Takagi et al., 2008***). Further, Kobayashi et al. showed the importance of TGF-β in a Th17 polarizing cytokine cocktail for the formation of CXCL13-producing Th cells (***Kobayashi et al., 2016***). Our findings show even more pronounced effects of FSK, with the FSK-mediated elevation of cAMP during Th17 differentiation inducing the skewing of the classic Th17 cell phenotype towards CXCL13-expressing Th cells.

In conclusion, the FSK-mediated induction of intracellular cAMP significantly modifies human Th cell differentiation *in vitro* by reinforcing the Th2 phenotype and by suppressing the Th17 phenotype. The specific modulation of cAMP represents a novel therapeutic target for the treatment of Th cell-related pathological processes.

## 4. Methods

### Isolation of Th naïve cells

Blood samples were collected from healthy volunteers of both sexes aged 18–60, which had been free of any medication and any signs of cold or other disease for at least 3 weeks before blood draw. They gave their informed consent based on the collection protocol approved by The Ethics Committee for Research at Masaryk University, Brno, Czech Republic (number EKV-2018-083) approved on 5th November 2018. The PBMCs were isolated from the buffy coat (500 ml of blood) using dextran separation followed by Histopaque 1077 (*Sigma-Aldrich, St. Louis, MO, USA*) separation and the lysis of remaining erythrocytes by water hypotonic lysis. Th naïve cells (CD4^+^CD25^−^CD45RA^+^) were sorted by BD FACSAria II (*BD Biosciences, Franklin Lakes, NJ, USA*) using conjugate antibodies anti-CD4-Pacific Blue, anti-CD25-PE/Cy7, and anti-CD45RA-PE (all *SONY Biotechnology, San Jose, CA, USA*).

### Activation, differentiation, and treatment

Each well of 6-well plates (*VWR, Radnor, PA, USA)* was coated with anti-CD3 antibody (1 μg/ml in PBS) (*ExBio, Prague, Czech Republic)* at 4°C overnight. Naïve CD4^+^ T cells were seeded (4–5 × 10^5^ cells/ml) in RPMI 1640 containing 10 % (v/v) low endotoxin fetal bovine serum (FBS, *BioTech, Prague, Czech Republic*) and 1% penicillin-streptomycin (*Gibco, Dublin, Ireland*) and pretreated with 10 ng/ml of soluble anti-CD28 antibody (*ExBio, Prague, Czech Republic*) for 30 min. Activated cells were incubated for 5 days in the presence of the following: IL-12 (25 ng/ml, *PeproTech, London, UK*) and neutralizing antibody anti–IL-4 (1 μg/ml, *eBioscience, San Diego, CA, USA*) for Th1 differentiation; IL-4 (10 ng/mL *PeproTech, London, UK*) and neutralizing antibody anti–IFN-γ (1 μg/ml, *eBioscience, San Diego, CA, USA*) for Th2 differentiation; and IL-6 (25 ng/ml), IL-23 (25 ng/ml), IL-1β (25 ng/ml), TGF-β (0.25 ng/ml, all *PeproTech, London, UK)* and neutralizing antibodies anti–IFN-γ (1 μg/ml, *eBioscience, San Diego, CA, USA*) and anti–IL-4 (5 μg/ml, *eBioscience, San Diego, CA, USA*) for Th17 differentiation. For inducing cAMP formation, receptor independent activator FSK (1 or 10 μM, *Sigma-Aldrich, St. Louis, MO, USA*) was added 20 h after activation and differentiation treatments, and treated cells were incubated for 5 days altogether. For the inhibition of ACs, the inhibitor ddAdo (100 μM, *MedChemExpress, NJ, USA*) was added 20 h after activation and differentiation treatment, or 30 min prior to FSK treatment.

### Gene expression analysis

Total RNA was isolated from cells using the Cell/Tissue RNA Kit (*CatchGene, New Taipei City, Taiwan*). Isolated RNA was reversely transcribed into cDNA using the Reverse Transcription Kit according to the manufacturer’s protocol (*GeneriBiotech, Hradec Kralove, Czech Republic*). Real-Time quantitative PCR reaction was performed in a QuantStudio5 (*Thermo Fisher Scientific, Waltham, MA, USA*) or LightCycler480 instrument using Universal Probe Library (UPL, *Roche, Basel, Switzerland*) and TaqMan primers and probes (*Thermo Fisher Scientific, Waltham, MA, USA*) using a LightCycler480 Probe Master (*Roche, Basel, Switzerland*) with the following program: an initial denaturation step at 95°C for 15 min, followed by 45 cycles of amplification (95°C for 15 s and 60°C for 1 min) and a final cooling step at 40°C for 1 min. Specific primers for transcription factors, signature genes, and PDEs (UPL, **Supplementary data 9A**) and particular AC isoforms (TaqMan, **Supplementary data 9B**) were used and data were normalized to RPL13a mRNA presented as 2^−Δct^. The primers were designed to detect all splicing variants for particular PDEs and ACs.

### Determination of AC activity

Th cell samples were tested for their ability to produce cAMP, determined by competitive immunoanalysis using Time-Resolved Fluorescence Resonance Energy Transfer (TR-FRET) as a detection system (Lance Ultra cAMP kit from *Perkin Elmer, Waltham, MA, USA*) according to the manufacture instructions. Cells were stimulated by FSK (in the concentration range 1-100 μM) for 30 min at RT in the presence of pan-PDE inhibitor IBMX (500 μM, *Sigma-Aldrich, St. Louis, MO, USA*). The fluorescence signal, which is inversely proportional to the overall amount of cAMP, was read immediately using a Spark M10 spectrophotometer (*Tecan, Zürich, Switzerland*) with the excitation fluorescence at 320 nm and emission fluorescence at 620nm and 665nm. The lag time was 150 μs and the integration time was 500 μs. The fluorescence signal, which is inversely proportional to the overall amount of cAMP, was normalized as a ratio between the experimental (665 nm) and reference (620 nm) values and inverted.

### Production of cytokines

The production of cytokines was detected in culture media after 5 days of incubation. Samples were centrifuged (300 x g, 5 min, 4°C) and the supernatants were frozen (−20°C). IL-2 and the effector cytokines IFN-γ, IL-13 and IL-17A were detected by IL-2 Human Uncoated ELISA Kit, IFN gamma Human Uncoated ELISA Kit, IL-13 Human Uncoated ELISA Kit, and IL-17A Human Uncoated ELISA Kit (all *Thermo Fisher Scientific, Waltham, MA, USA*), respectively, according to the manufacturer’s instructions, using a Sunrise microplate reader (*Tecan, Zürich, Switzerland*).

### Single cell RNA sequencing

After 5 days of differentiation treatment, cells were collected with the aim of obtaining a single-cell suspension with high cell viability. Next, half a million cells per sample were incubated for 10 min in 50 μl of PBS with 3% FSB and 5 μl of blocking buffer (*BioLegend, San Diego, CA, USA*) and then stained with 50x diluted TotalSeq-B cell feature barcode antibody (*BioLegend, San Diego, CA, USA*, **Supplementary data 10A**) to barcode each sample for multiplexing for the next 30 min. Stained cells were then washed and resuspended in PBS with 3% FBS three times. Cells were resuspended in PBS buffer and recounted using Burker chamber counting. Cells from each sample were pooled in equal cell numbers.

10 000 cells were loaded into the well of a 10×-Genomics Chromium microfluidic chip to create a GEM (a gel bead in emulsion). Upon encapsulation and reverse transcription, the cDNA was processed for single-cell RNA-sequencing using the Chromium Next GEM Single Cell 3′ Reagent Kits v3.1 Gene Expression Library With Feature Barcoding technology for Cell Surface Protein (*10×-Genomics, Pleasanton, CA, USA*), according to the manufacturer’s instructions. Libraries were sequenced on an Illumina NextSeq 500/550 system using NextSeq 500/550 High Output Kit v2.5 (75-cycles, *Illumina, San Diego, CA, USA*).

Single-cell RNA-sequencing data were pre-processed using the standardized protocol from the Cell Ranger Single-Cell Software Suite (*v3.1.0; 10X Genomics, Pleasanton, CA, USA*). Upon the first quality control, particular reads were assigned to cells and then aligned to the reference genome using the hg38 assembly of the human genome (*GRCh38-3.0.0*). Subsequently, gene expression was quantified using the UMI counts located in specific cell barcode sequences. The pre-processed data from Cell Ranger were imported into RStudio (*v4.0.4, RStudio, PBC, Boston, MA, USA*) and analyzed using the Seurat package (***Hao et al., 2021***) (*v4.0.0*). Firstly, a check for the presence of Antibody Capture and reads was performed. Cell Feature Barcodes were then renamed, and to each of them, we assigned a sample label corresponding to our experiment setup. The Seurat object was created using the two filtration parameters (**Supplementary data 10**.). First, all genes expressed in less than 3 cells were excluded from the analysis (**Supplementary data 10**.). The second step was the filtration of cells with less than 200 genes expressed to exclude dead cells or empty droplets. The HTO assay was normalized using the default normalization method. Upon demultiplexing with a 0.99 positive quantile parameter set, negative cells and doublets were removed. From the overall 10 804 cells that were captured (after the pre-processing step), 8 941 cells were classified as singlets and used for further analysis (**Supplementary data 10**.). Each of the particular steps was followed by a quality control check. The median values are presented **in Supplementary data 10**. Later on, we further filtrated our dataset by excluding all cells with more than 20% of mitochondrial genes present in the sequenced transcriptome (**Supplementary data 10A**). Our dataset was simultaneously filtrated regarding the number of genes present in each cell (**Supplementary data 10B**). As mentioned earlier, the filtration excluded another 1 799 potentially non-viable cells from our analysis (the number of singlet cells for each sample is illustrated in **Table 1**.). In total, the percentage of viable Singlet cells was 66.1% of the raw cell count.

**Table 1.** Amount of cells in each sample used for data analysis after filtration with respect to nFeature_RNA and the % of mitochondrial gene expression threshold

| Sample ID | NT ctrl | ACT | ACT + FSK | Th1 | Th1 + FSK | Th2 | Th2 + FSK | Th17 | Th17 + FSK | TOTAL |
| --- | --- | --- | --- | --- | --- | --- | --- | --- | --- | --- |
| No. of cells | 781 | 961 | 745 | 785 | 840 | 912 | 666 | 724 | 728 | 7142 |

On such a filtrated dataset, we ran the *FindVariableFeatures* function with default parameters to identify outliers on a mean variability plot. The ‘number of genes’ threshold was set as close as possible to the observed median values (**Supplementary data 10**). In the following step of the analysis, we reduced our dataset to only protein-coding genes, using the Biostrings-based genome data package (***Pagès, 2021***) (*v1.58.0*) and genome annotated sequences for Homo sapiens (BSgenome.Hsapiens.NCBI.GRCh38) provided by National Center for Biotechnology Information (NCBI, GenBank accession number: GCA_000001405.28; 2019-02-28).

Subsequently, the *SCTransform* (***Hafemeister and Satija, 2019***) package was used to transform and scale the data. Principal component analysis (PCA) with default parameters was run on transformed data to reduce dimensionality. On the basis of the PCA, uniform manifold approximation and projection for dimension reduction (UMAP (***Becht et al., 2018***)) were performed (Supplementary data 10). This step was followed by SNN (shared nearest neighbours) clustering. We identified clusters (with a resolution threshold equal to 0.9) according to transcription profile similarities.

Using the *FindMarkers* function, we identified the most differentially expressed genes in each cluster, and therefore, were able to distinguish artificially formed clusters from clusters with biological relevance to our dataset. Various treatment types (subsets) were also extracted from the dataset on the basis of sample labels. Extracted sub-datasets (Th1 & Th1+FSK, Th2 & Th2+FSK, Th17 & Th17+FSK) underwent differential expression analysis using the Model-based Analysis of Single Cell Transcriptomics package, (*MAST, v1.16.0, (**Finak et al., 2015**)*). Respectively, a list of the 50 most differentially expressed genes was used to perform gene enrichment analysis using the *enrichR* package (*v3.0*, (***Kuleshov et al., 2016***)). Genes used as inputs were sorted into categories according to annotations in the GO_Molecular_Function_2018 library.

In addition, specific sub-datasets were extracted using automatic cluster assembly. Clusters were extracted again and used for differential expression analysis based on the MAST algorithm and the sample labels. In accord with the literature, such extracted sub-datasets were used to visualize signature genes specific to distinct treatment types.

### Western Blot Analysis

Protein expression analysis of target proteins was performed as described previously (***Mosejova et al., 2021***). Cells were lysed using lysis buffer (1% sodium dodecyl sulfate (SDS); 100 mM Tris,pH 7.4; 10% glycerol). The protein concentration was determined using BCA™ protein assay (*Pierce, Rockford, IL, USA*) using BSA (*Sigma- Aldrich, St. Louis, MO, USA*) as a standard, and 10-20 μg/well of total protein supplemented with Laemli buffer (200 mM Tris-HCl, pH 6.8; 3% SDS; 30% glycerol; 0.03 bromphenol blue; 3% ß-mercaptoethanol; 200 mM dithiothreitol) were run in SDS-polyacrylamide gel electrophoresis, using 10% running gel. Separated proteins were transferred to polyvinylidenedifluoride membranes that were blocked in 5% non-fat milk or 5% BSA (*Sigma-Aldrich, St. Louis, MO, USA*) in Tris-buffered saline with 0.1% Tween 20 (TBS-T). Then, the membranes were incubated with antibodies against Vinculin (E1E9V) XP^®^ Rabbit mAb (1:1000; #13901, *Cell signaling technologies, Danvers, MA, USA*), Sestrin2 (1:1000; #8487S, *Cell signaling technologies, Danvers, MA, USA*), and PPAR-γ (1:1000; #2435S, *Cell signaling technologies, Danvers, MA, USA*) in 5% non-fat milk/TBS-T (vinculin) or 5% BSA/TBS-T (Sestrin2, PPAR-γ) at 4°C overnight. Corresponding secondary HRP-conjugated anti-rabbit (1:3000 or 1:2000 in 5% non-fat milk or 5% BSA/TBS-T; *Sigma-Aldrich, St. Louis, MO, USA*) was used for a 1-hour incubation at room temperature. The luminescence signal was read on an Amersham Imager680 (*GE Healthcare, North Richland Hills, TX, USA*). The relative levels of proteins were quantified by scanning densitometry using the ImageJ™ program v1.48 (*National Institutes of Health, Bethesda, MD, USA*).

### Measurement of Metabolic Phenotype and Potential by Seahorse Assay

The metabolic phenotype was measured by an Agilent Seahorse XFp extracellular flux analyzer (*Agilent, Santa Clara, CA, USA*) according to the manufacturer’s protocol. The Agilent Seahorse XF Cell Energy Phenotype Test (*Agilent, Santa Clara, CA, USA*) allows simultaneous measurement of the two major energy-producing pathways - mitochondrial respiration and glycolysis – under baseline and stressed conditions, giving information about baseline phenotype, stressed phenotype, and metabolic potential. Selected samples underwent the Th2 differentiation process as well as FSK (10 μM) treatment, as described in the “Activation, differentiation and treatment” chapter of the Methods section. On day 5, cells were collected and washed twice in freshly prepared Agilent Seahorse XF Base Medium with 1 mM pyruvate, 2 mM glutamine, and 10 mM glucose (assay medium). An Agilent Seahorse XFp Sensor Cartridge was hydrated in Agilent Seahorse XF Calibrant at 37 °C in a non-CO2 incubator overnight. Next, cells in a concentration of 2×10^5^ cells/well were transferred into an 8-well plate (Agilent Seahorse XFp miniplate), which was previously coated overnight with poly-L-lysine (*Santa Cruz Biotechnology, Dallas, TX, USA)* at 8°C and, prior to cell seeding, washed with H_2_O and air-dried. Cells in the plate were incubated in a non-CO2 incubator for 1 h. The stressed condition was induced using carbonyl cyanide-p-trifluoromethoxyphenylhydrazone (FCCP) and oligomycin (both 1 μM, *MedChemExpress, NJ, USA*). Oligomycin, as an inhibitor of ATP synthase, causes a compensatory increase in the rate of glycolysis, as the cells attempt to meet their energy demands via the glycolytic pathway, while FCCP depolarizes the mitochondrial membrane and drives oxygen consumption rates higher, as the mitochondria attempt to restore the mitochondrial membrane potential. Agilent Wave software was used for data evaluation.

### Flow cytometric analysis – intracellular staining

Cells were cultured for 5 h with 4 μM monensin (*Sigma-Aldrich, St. Louis, MO, USA*, protein transport inhibitor). The intracellular molecule CXCL13 was stained with CXCL13-PE (*Thermo Fisher Scientific, Waltham, MA, USA*) using Foxp3/Transcription Factor Staining Buffer Set (*eBioscience, San Diego, CA, USA*). Analysis was performed using a SONY SP6800 Spectral Analyzer (*SONY Biotechnology, San Jose, CA, USA*) and data were analyzed with FlowJo Version 10 (*Tree Star, Ashland, OR, USA*). Fixable Viability Dye eFluor 506 (*eBioscience, San Diego, CA, USA*) was used to exclude dead cells. The borders of gates were determined according to unstained control. Data are expressed as geometric mean of fluorescence.

### Statistics

Data are presented as mean ± standard error of the mean (SEM). Statistical differences between mean values were determined using GraphPad Prism-6 software (*GraphPad software, La Jolla, CA, USA*) and tested by one-way ANOVA or unpaired two-tailed Student’s t-test. One-sample t-test was used to compare values expressed as percentages. In the case of one sample t-test, Bonferroni correction of the p value for multiple-comparisons was performed. The number of independent repeats (n) is given in each figure legend.

## Institutional Review Board Statement

The study was conducted according to the guidelines of the Declaration of Helsinki, and approved by the Institutional Ethics Committee of Masaryk University, Brno, Czech Republic (number EKV-2018-083).

## Informed Consent Statement

Informed consent was obtained from all subjects involved in the study.

## Data Availability Statement

The data presented in this study are available on request from the corresponding author.

## Acknowledgement

The authors are grateful to Natalia Vadovicova for her help with the determination of cell metabolic activity, Radek Fedr who provided technical support for flow cytometry, and Terézia Kurucová, Petr Tauš, Karla Plevová, and Boris Tichý for the help with single cell RNA sequencing experiments.

