## Supplementary data for "A forskolin-mediated increase in cAMP promotes T helper cell differentiation into the Th1 and Th2 subsets rather than into the Th17 subset"

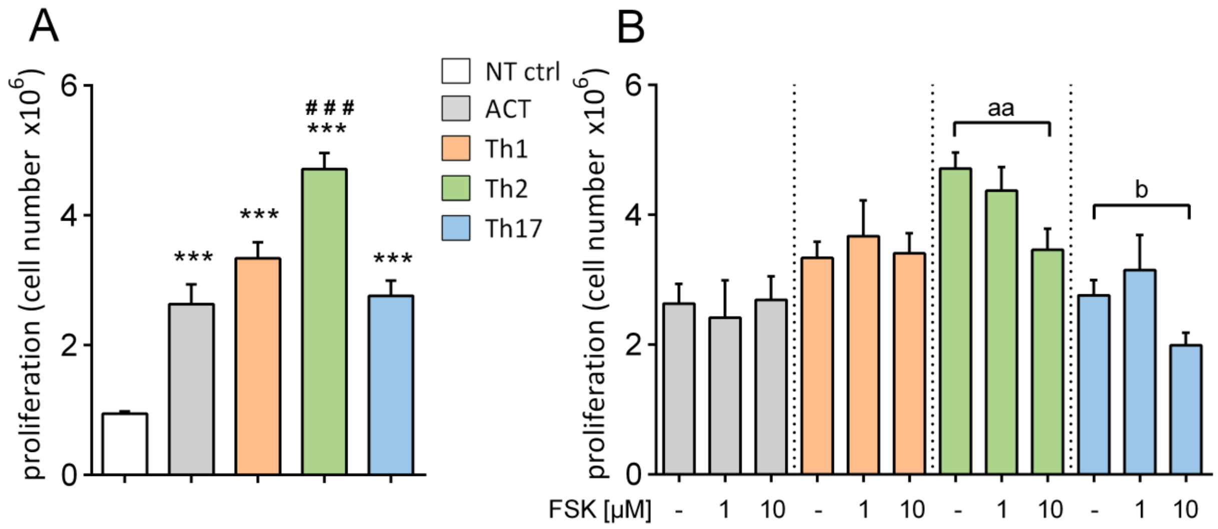


***Supplementary data 1. A.*** *Proliferation of the Th1, Th2, and Th17 subset upon 5-day TCR activation (anti-CD3, anti-CD28) and/or differentiation (specific cytokine and inhibitory antibody cocktails) compared to ACT (activated control) and NT ctrl (non-treated control, resting cells).* ***B.*** *Comparison of ACT, Th1, Th2, and Th17 cell proliferation with T1, Th2, or Th17 cells treated with FSK (1 or 10 µM). Statistical significance was examined by one-away ANOVA (* vs NT ctrl, # vs ACT) or unpaired two-tailed Student's t-test (a vs Th2, b vs Th17). Mean ± SEM, n=7-15,*$p < 0.05$*, ***$p < 0.01$*, ****$p < 0.001$

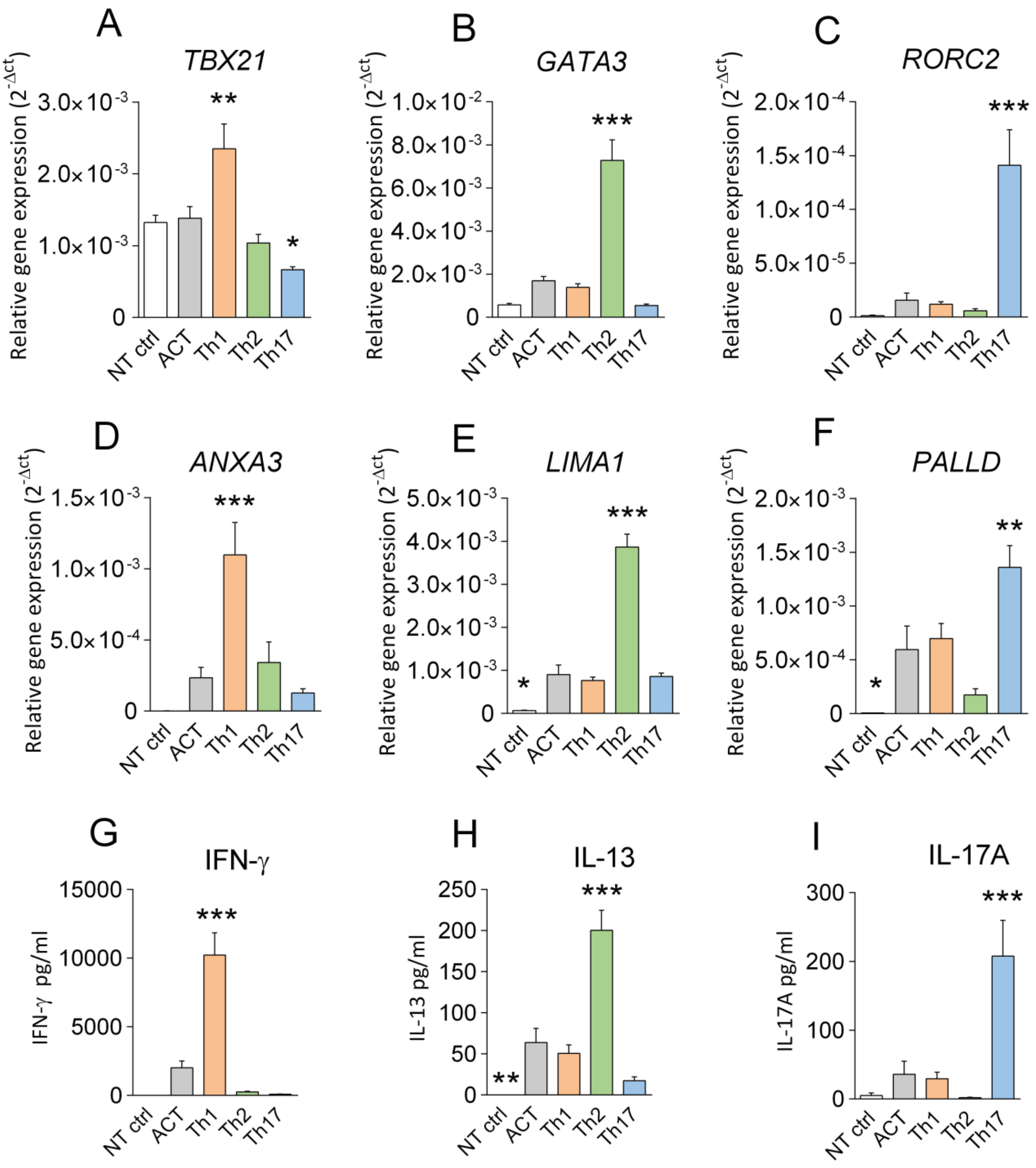


***Supplementary data 2.*** *Verification of 5-day differentiation into Th1, Th2, and Th17 subsets.* ***A-F.*** *Relative gene expression analysis of specific markers for Th1 (TBX21 and ANXA3), Th2 (GATA3 and LIMA1) and Th17 (RORC2 and PALLD).* ***G-I.*** *Production of effector cytokines IFN-γ (Th1), IL-13 (Th2) and IL-17A (Th17) in Th1, Th2, and Th17 subsets. ACT – activated control, NT ctrl – non-treated control (resting cells). Statistical significance was examined by one-way ANOVA (* vs ACT). Mean ± SEM, n=9-13, *p < 0.05, **p < 0.01, ***p < 0.001.*

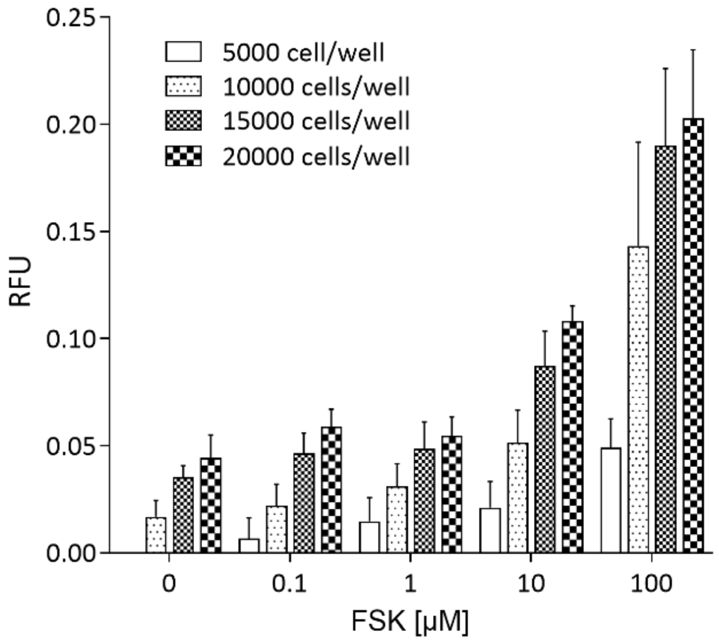


***Supplementary data 3.*** *TR-FRET detection of FSK-mediated cAMP production in T helper naive cells. 5000, 10000, 15000 and 20000 cells/well and concentration range 0.1-100 µM FSK were used for 30 min stimulation.*

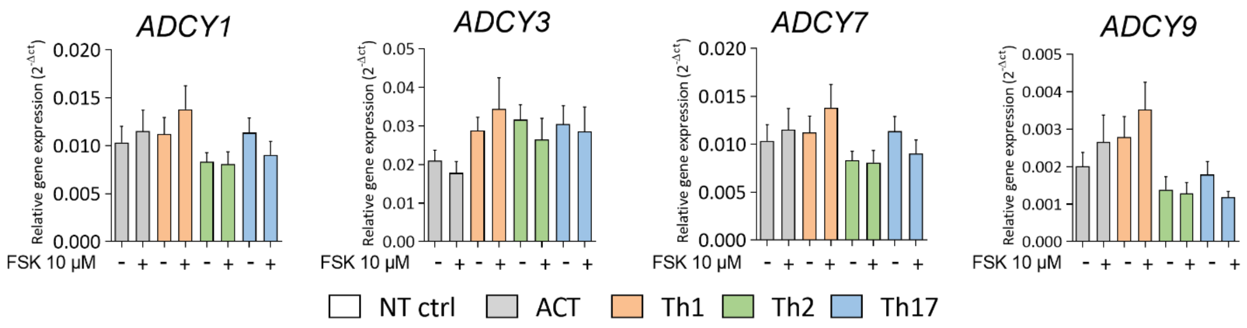


***Supplementary data 4.*** *Relative gene expression analysis of AC isoforms ADCY1, ADCY3, ADCY7 and ADCY9 in Th1, Th2 and Th17 cells, and upon FSK (10 µM) treatment. Mean ± SEM, n=9-13*

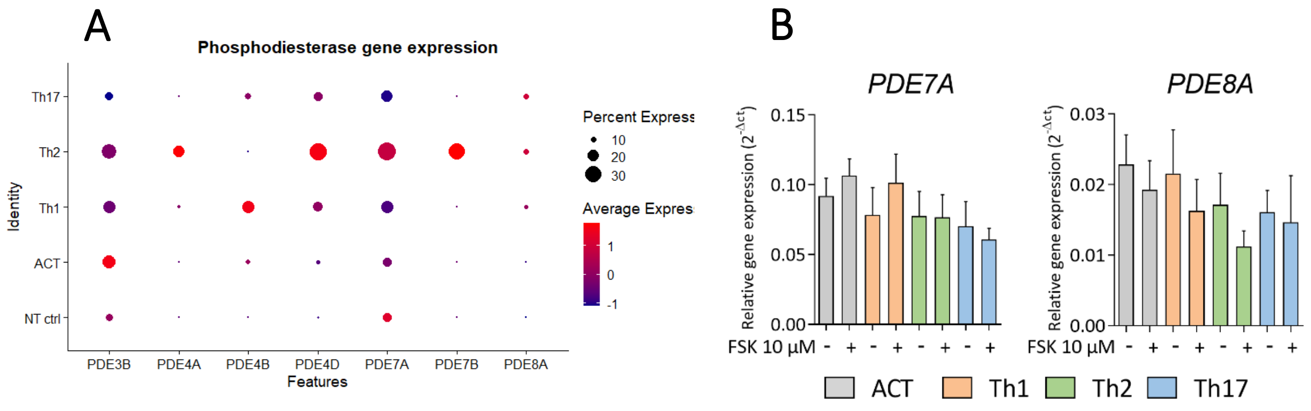


***Supplementary data 5.*** *Relative gene expression analysis of PDEs in Th1, Th2, and Th17 cells.* ***A****. scRNAseq DotPlot visualization of detectable PDE isoforms in NT ctrl, ACT, Th1, Th2, and Th17 cells****.*** *Different colors correspond to the scaled expression (Z-score); the dot size represents the percentage of positive cells.* ***B.*** *Gene expression analysis of PDE isoforms PDE7A and PDE8A in Th1, Th2 and Th17 cells, and upon FSK (10 µM) treatment. ACT – activated, NT ctrl – non-treated control. Mean ± SEM, n=8.*

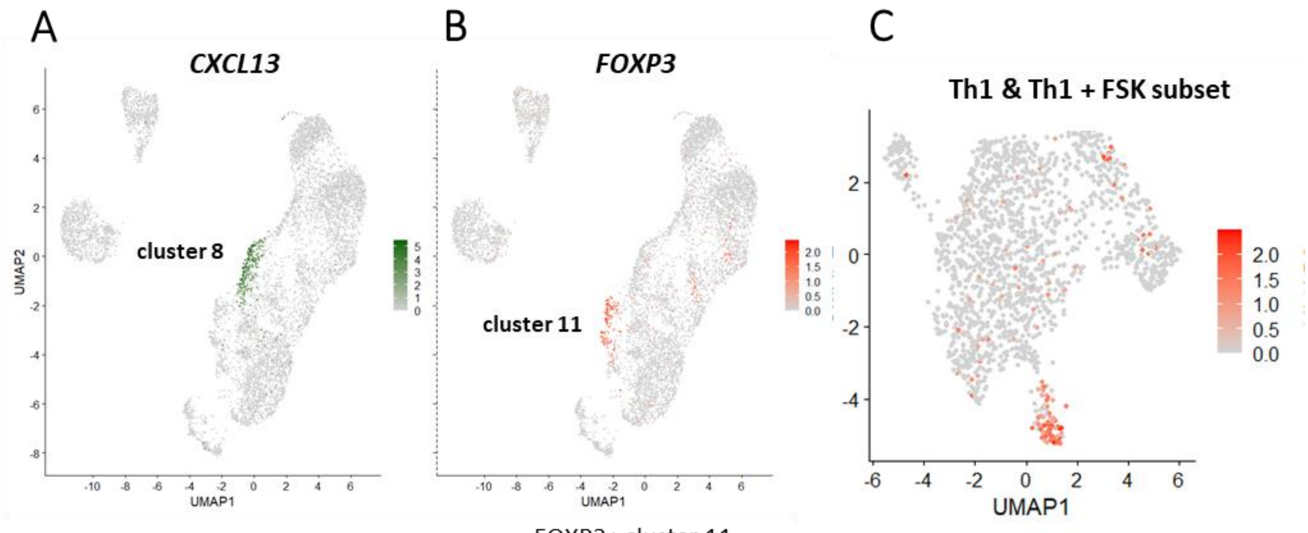


CXCL13+ cluster 8

| Sample ID | No. of cells | % |
|---|---|---|
| NT ctrl | 0 | 0,0 |
| ACT | 0 | 0,0 |
| ACT + FSK | 4 | 1,2 |
| Th1 | 0 | 0,0 |
| Th1 + FSK | 0 | 0,0 |
| Th2 | 0 | 0,0 |
| Th2 + FSK | 0 | 0,0 |
| Th17 | 7 | 2,1 |
| Th17 + FSK | 322 | 96,7 |
| **TOTAL** | **333** | **100** |

FOXP3+ cluster 11

| Sample ID | No. of cells | % |
|---|---|---|
| NT ctrl | 0 | 0,0 |
| ACT | 21 | 8,5 |
| ACT + FSK | 18 | 7,3 |
| Th1 | 44 | 17,9 |
| Th1 + FSK | 68 | 27,6 |
| Th2 | 15 | 6,1 |
| Th2 + FSK | 20 | 8,1 |
| Th17 | 30 | 12,2 |
| Th17 + FSK | 30 | 12,2 |
| **TOTAL** | **246** | **100** |

(Th1 and Th1 + FSK: 45,5%)

***Supplementary data 6. A,B.*** *UMAP visualisation of CXCL13+ and FOXP3+ cells, with tables of numbers and percentages of cell subsets creating cluster 8 or 11.* ***C.*** *UMAP visualisation of FOXP3 in extracted dataset Th1 and Th1+FSK.*

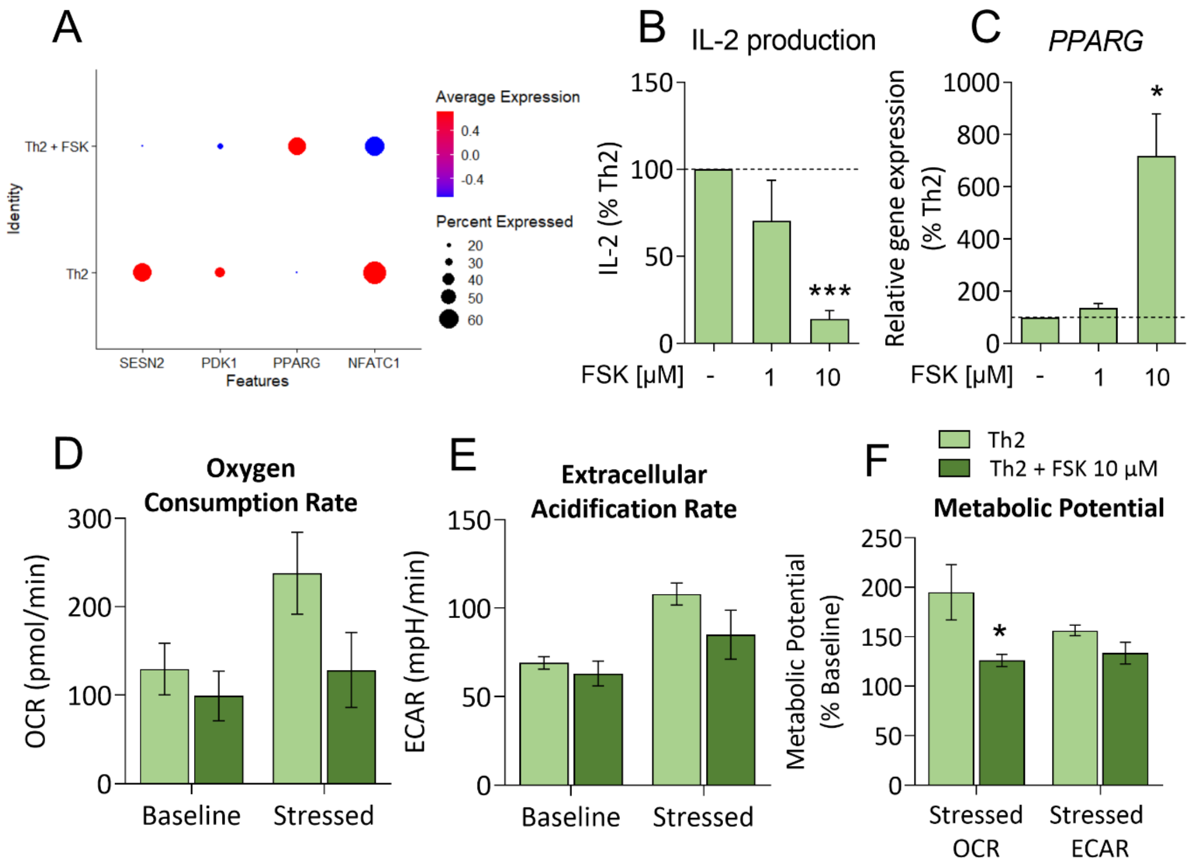


***Supplementary data 7. A.*** *DotPlot visualization of scRNAseq analysis of SESN2, PDK1, PPARG and NFATC1 expressions in Th2 cells and Th2 cells upon treatment with 10 μM FSK (Th2+FSK). Different colors correspond to the scaled expression (Z-score), the dot size represents the percentage of positive cells.* ***B,C.*** *Analysis of IL-2 production and relative gene expresion of PPARG in Th2 cells and Th2 cells upon FSK treatment (1 or 10 μM). Statistical significance was examined by one-sample t-test, separately comparing the effect of each experimental treatment with Th1 cells. Mean ± SEM, n=3-7, *p < 0.05, ***p < 0.001.* ***D-F.*** *Analysis of mitochondrial respiration (OCR, oxygen consumption rate) and glycolysis (ECAR, extracellular acidification rate) using Seahorse XF Cell Energy Phenotype test. Visualisation of OCR, ECAR and metabolic potential under the baseline or stressed condition induced with stressor compounds FCCP and oligomycin (both 1 μM). Statistical significance was examined by unpaired two-tailed Student's t-test. Mean ± SEM, n=4, *p < 0.05*

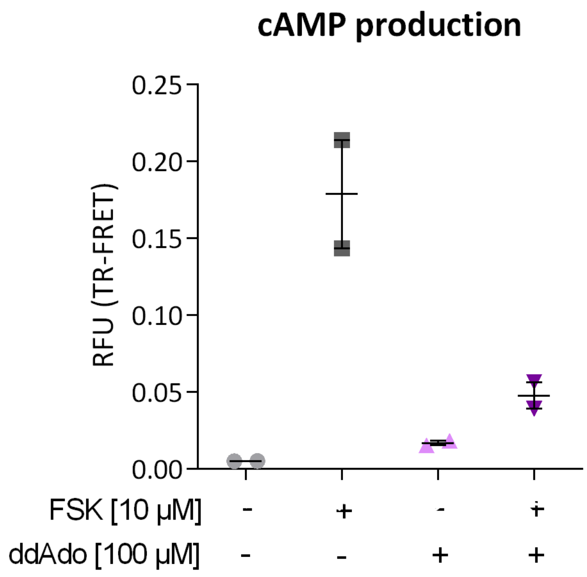


***Supplementary data 8.*** *TR-FRET detection of cAMP production in T helper cells activated for 20 h. 20000 cells/well were treated for 30 min with 10 µM FSK, 100 µM ddAdo or with combination of the two compounds.*

| Gene of interest | Forward primer (5´→ 3´) Reverse primer (5´→ 3´) | Amplicon length | UPL probe |
|---|---|---|---|
| *RPL13A* | cattgttgccctggaatgta<br>ccttgctcccagcttcctat | 89 nt | #27 |
| *TBX21* | cccaactgtcaattccttgg<br>gggaacaggatactggttgg | 66 nt | #55 |
| *GATA3* | tcagaccaccacaaccacac<br>tcttcatagtcaggggtctgttaat | 113 nt | #25 |
| *RORC2* | cagcgctccaacatcttct<br>cacaccgttcccacatctc | 79 nt | #6 |
| *PDE3B* | aacaatggtataagcctcattatcaa<br>cgagcctcatttagcactga | 76 nt | #10 |
| *PDE4A* | gccacgcctgcactagat<br>ccagggtgatccacatcg | 89 nt | #85 |
| *PDE4B* | caacactgaaaatgaagatcacct<br>tgcatgttaggggtctattgtg | 110 nt | #5 |
| *PDE4D* | tgcttccgtggaaaaatca<br>caaaatatcctgggcgtcag | 103 nt | #14 |
| *PDE7A* | ctgcattatacattcgtatgctagg<br>tgagagtggaaaatacgaaaatca | 112 nt | #15 |
| *PDE7B* | caggacaggcactttatgctt<br>cctttcactccactgcttgc | 96 nt | #14 |
| *PDE8A* | tcatcatagaccacagaaatcctc<br>agagttttgatgatctgatagacctg | 75 nt | #42 |
| *ANXA3* | cagaaatatcagccaaaaggacat<br>ggcgtgttcctcacacaat | 105 nt | #56 |
| *LIMA1* | gcaagtggaaggaagatctctg<br>ctccgagtcaaatgtagaagatga | 93 nt | #85 |
| *PALLD* | tgatcatagagccagtcacgtc<br>gggggtttgtgtgcttctt | 124 nt | #15 |
| *CXCL13* | gcaggcttctatgaaagactcaa<br>ctcttgacaggctcaagttcc | 65 nt | #16 |
| *PPARG* | gacaggaaagacaacagacaaatc<br>ggggtgatgtgtttgaacttg | 96 nt | #7 |
| *SESN2* | ccatcacccgctacatgacc<br>gagaagtccttgtggggagc | 71 nt | #15 |

| Gene symbol | Assay ID |
|---|---|
| ADCY1 | Hs00299832_m1 |
| ADCY2 | Hs01058848_m1 |
| ADCY3 | Hs01086502_m1 |
| ADCY4 | Hs00934104_m1 |
| ADCY5 | Hs02890018_m1 |
| ADCY6 | Hs00209600_m1 |
| ADCY7 | Hs00181579_m1 |
| ADCY8 | Hs02890018_m1 |
| ADCY9 | Hs00181599_m1 |
| ADCY10 | Hs01037968_m1 |

***Supplementary data 9.*** *List of the primers used in quantitative RT-PCR.* ***A.*** *Sequence and probe numbers of UPL primers.* ***B.*** *List of the TaqMan primers*

A

| Sample ID | # | Product ID | Barcode sequence |
|---|---|---|---|
| NT ctrl | 1 | TotalSeq-B0251 anti-human Hashtag 1 | GTCAACTCTTTAGCG |
| ACT | 2 | TotalSeq-B0252 anti-human Hashtag 2 | TGATGGCCTATTGGG |
| ACT + FSK | 3 | TotalSeq-B0253 anti-human Hashtag 3 | TTCCGCCTCTCTTTG |
| Th1 | 4 | TotalSeq-B0254 anti-human Hashtag 4 | AGTAAGTTCAGCGTA |
| Th1 + FSK | 5 | TotalSeq-B0255 anti-human Hashtag 5 | AAGTATCGTTTCGCA |
| Th2 | 6 | TotalSeq-B0256 anti-human Hashtag 6 | GGTTGCCAGATGTCA |
| Th2 + FSK | 7 | TotalSeq-B0257 anti-human Hashtag 7 | TGTCTTTCCTGCCAG |
| Th17 | 8 | TotalSeq-B0258 anti-human Hashtag 8 | CTCCTCTGCAATTAC |
| Th17 + FSK | 9 | TotalSeq-B0259 anti-human Hashtag 9 | CAGTAGTCACGGTCA |

B

| Cell ranger Pre-processing | |
|---|---|
| Coverage (%) | 97.7 |
| UMI counts median | 8 658 |
| Median genes per cell | 2649 |
| Median reads per cell | 21 376 |

| Cell demultiplexing | |
|---|---|
| Doublets | 505 |
| Negative | 1358 |
| Singlets | 8941 |

| Singlet median values | |
|---|---|
| Transcripts | 6582 |
| Gene count | 1849 |

| Seurat analysis parameters | |
|---|---|
| HTO assay method | CLR |
| Cell dropout number of genes | < 3 |
| Gene threshold for singlets | > 200 |
| Variable features | 2000 |
| SCTransforms (genes/cells) | 3000/7000 |
| UMAP number of dimensions | 50 |
| UMAP cluster resolution | 0.7;0.9 |

C

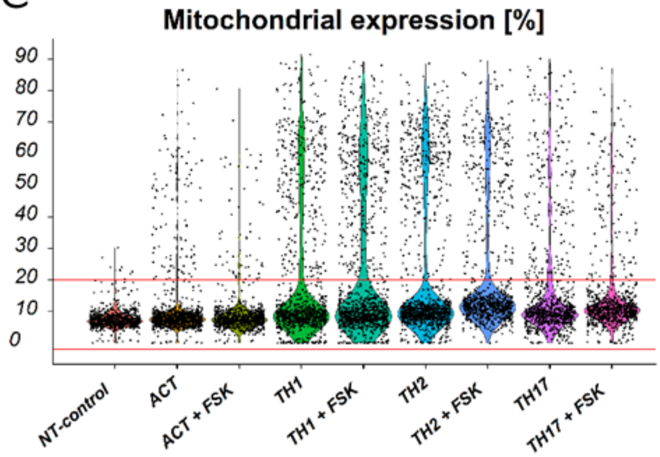


D

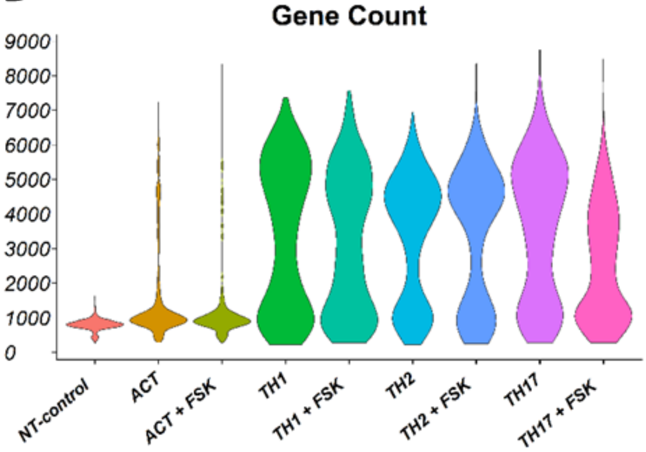


***Supplementary data 10. A.*** *Sequences of TotalSeq-B hashtag antibodies used for labelling of particular samples.* ***B.*** *The table of chosen parameters for scRNAseq analysis pipeline.* ***C.*** *Quality control of percentage of mitochondrial gene expression among samples. The Red line corresponds to the threshold set for further analysis.* ***D.*** *Violin plot of a number of gene abundance in each sample. NT-control, ACT, and ACT + FSK are transcriptionally attenuated.*
